# ADR-2 regulates fertility and oocyte fate in *C. elegans*

**DOI:** 10.1101/2023.11.01.565157

**Authors:** Emily A. Erdmann, Melanie Forbes, Margaret Becker, Sarina Perez, Heather A. Hundley

## Abstract

RNA binding proteins play essential roles in coordinating germline gene expression and development in all organisms. Here, we report that loss of ADR-2, a member of the Adenosine DeAminase acting on RNA (ADAR) family of RNA binding proteins and the sole adenosine-to-inosine RNA editing enzyme in *C. elegans*, can improve fertility in multiple genetic backgrounds. First, we show that loss of RNA editing by ADR-2 restores normal embryo production to subfertile animals that transgenically express a vitellogenin (yolk protein) fusion to green fluorescent protein. Using this phenotype, a high-throughput screen was designed to identify RNA binding proteins that when depleted yield synthetic phenotypes with loss of *adr-2*. The screen uncovered a genetic interaction between ADR-2 and SQD-1, a member of the heterogenous nuclear ribonucleoprotein (hnRNP) family of RNA binding proteins. Microscopy, reproductive assays, and high-throughput sequencing reveal that *sqd-1* is essential for the onset of oogenesis and oogenic gene expression in young adult animals, and that loss of *adr-2* can counteract the effects of loss of *sqd-1* on gene expression and rescue the switch from spermatogenesis to oogenesis. Together, these data demonstrate that ADR-2 can contribute to the suppression of fertility and suggest novel roles for both RNA editing-dependent and independent mechanisms in regulating embryogenesis.

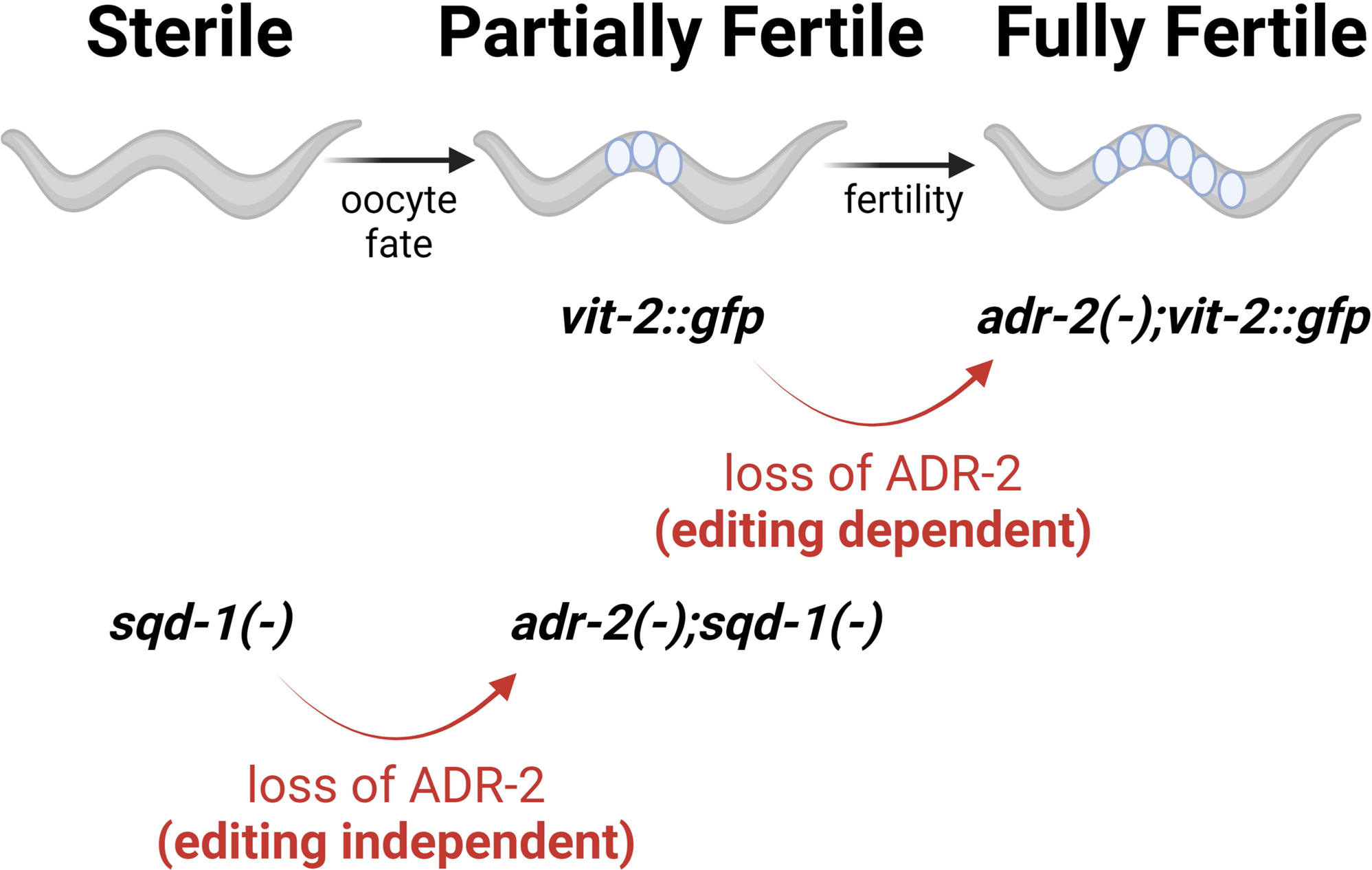
Graphical Abstract: Figure created with BioRender.

**Article Summary:** The molecular information that controls early development is RNA. Interactions between RNA and RNA binding proteins (RBPs) are critical for successful reproduction across species. In this study, we uncovered two roles for the RBP ADR-2 in regulating reproduction. First, we demonstrate that the RNA editing activity of ADR-2 regulates fertility. Next, we screened over 250 other RBPs revealed a genetic interaction between ADR-2 and SQD-1. Further analysis revealed that *sqd-1* is essential for the onset of oogenesis, and an editing-independent function of ADR-2 influences proper germline gene expression and oocyte fate in these animals.

## Introduction

Underlying successful reproduction is a complex gene regulatory network that coordinates many cellular and molecular events within the germline to ensure proper reproductive timing, development of gametes, and embryogenesis. Because many of these processes require rapid responses to signals or transient gene expression changes during times of transcriptional quiescence, RNA-level regulation is highly relied upon in germ cells and supporting tissues to coordinate gene expression (Seydoux and Braun 2006). As such, many RNA binding proteins (RBPs) have been found to be highly expressed during gametogenesis and essential for proper germline function and reproduction (Albarqi and Ryder 2022; Nguyen-Chi and Morello 2011; Rosario and others 2017). However, as fertility is essential for the viability of a species, there are also numerous epistatic and compensatory interactions that can obscure the identification of important factors in RNA regulation in germ cells and supporting cells.

*Caenorhabditis elegans* is a useful model organism for germline studies because these animals exist mainly as hermaphrodites, producing both sperm and oocytes at different life stages. Studies of the *C. elegans* germline have identified RBPs that regulate stability and translation of specific transcripts to coordinate various reproductive processes, including germline stem cell renewal (Subramaniam and Seydoux 2003; Wang and others 2020), the transition from spermatogenesis to oogenesis (Bachorik and Kimble 2005; Datla and others 2014; Kim and others 2012; Yoon and others 2017; Zanetti and others 2012) and from mitosis to meiosis (Kimble and Crittenden 2007; Park and others 2020; Priti and Subramaniam 2015) as well as proper progression through meiosis (Albarqi and Ryder 2022; Spike and others 2014).

One of the final steps in *C. elegans* oogenesis is the provisioning of yolk into maturing oocytes, which is also known to be regulated by at least one RBP (Lee and Schedl 2001). While viable embryos can be formed in the absence of yolk (Van Rompay and others 2015), yolk contributes to the fitness of progeny (Dowen and Ahmed 2019; Jordan and others 2019; Kern and others 2021). A major component of yolk is vitellogenin, or yolk protein. Vitellogenins are synthesized in the adult intestine, then exported into the body cavity and taken up into developing oocytes via receptor-mediated endocytosis (RME) (Kimble and Sharrock 1983). In a previous study, we observed that animals expressing a transgenic fusion of the vitellogenin gene *vit-2* with green fluorescent protein (*gfp*) contain fewer embryos and produce fewer progeny than wild-type animals (Erdmann and others 2022).

Here, we describe our finding that the fertility defects of *vit-2::gfp* animals are rescued in animals lacking *adr-2*, a member of the Adenosine DeAminase acting on RNA (ADAR) family of RNA binding proteins. ADARs bind double-stranded RNA and can affect gene expression either through binding transcripts or by catalyzing the deamination of adenosine (A) to inosine (I), a process known as A-to-I RNA editing (Erdmann and others 2021; Goldeck and others 2022; Savva and others 2012). ADR-2 is the sole A-to-I RNA editing enzyme in *C. elegans* (Arribere and others 2020), and has been shown to play important roles in chemotaxis (Deffit and others 2017; Tonkin and others 2002) as well as resistance to pathogen infection (Dhakal and others 2024) and hypoxia (Mahapatra and others 2023).

The ADR-2 dependent fertility defects in *vit-2::gfp* animals reported here, along with previously reported synthetic germline phenotypes of ADARs (Fischer and Ruvkun 2020; Reich and others 2018), prompted further exploration of the role of ADR-2 in germline RNA regulatory pathways. As such, we designed a high-throughput screen to identify synthetic germline phenotypes between ADR-2 and other RBPs. The screen revealed a genetic interaction between *adr-2* and *sqd-1*, an RBP orthologous to *Drosophila melanogaster* Squid and *Homo sapiens* hnRNPs (heterogeneous nuclear ribonucleoproteins) (Kim and others 2018; Shaye and Greenwald 2011). While *sqd-1* has not been well-characterized, a genome-wide RNA interference (RNAi) screen previously reported that decreased *sqd-1* expression impacted fertility in *C. elegans* (Maeda and others 2001). In addition, SQD-1 has also been reported to physically interact with GLD-1 (Akay and others 2013), an RBP critical to regulate RNA patterning and translational repression in the germline (Scheckel and others 2012). *Drosophila squid* is known to direct embryo patterning via RNA localization (Clouse and others 2008; Goodrich and others 2004; Kelley 1993; Lall and others 1999; Norvell and others 1999) and has recently been shown to be required for germline stem cell maintenance (Finger and others 2023). Here, we report the first examination of the effects of depleting *sqd-1* on germline morphology and gene expression. We show that germlines depleted of *sqd-1* fail to initiate oogenesis. Additionally, we show that loss of *adr-2* partially restores oogenesis and oogenic gene expression in animals lacking *sqd-1*. Furthermore, we demonstrate that both ADR-2 and SQD-1 are expressed throughout undifferentiated germ cells and oocytes, but are not present in sperm, suggesting expression of these factors is specific to oocytes and may contribute to specifying oocyte fate.

Together, the data presented here show that both RNA editing and editing-independent functions of ADR-2 can regulate fertility in *C. elegans* and suggest a broader role for ADR-2 in contributing to germline RNA regulation to achieve optimal reproductive function.

## Materials and Methods

### Worm strains and maintenance

All worm strains were maintained at 20⁰C on nematode growth media (NGM) seeded with *Escherichia coli* OP50. Worms were thawed regularly from frozen stocks to minimize effects of accumulated random mutations. The following previously generated strains were used in this study: Bristol Strain N2, BB20 (*adr-2(ok735)*) and BB21 (*adr-1(tm668);adr-2(ok735)*) (Hundley and others 2008), VC1106 (*sqd-1(ok1582) IV/nT1[qIs51](IV;V)*) (CGC), HAH20 (*pwIs23[vit-2::gfp]*) (Erdmann and others 2022), HAH10 (*adr-2(G184R)*) (Deffit and others 2017), *adr-2(uu28)* (a kind gift from Brenda Bass), HAH58 (3XFLAG::ADR-2) (Dhakal and others 2024). Strains generated in this study include: HAH51 (*adr-2(ok735);pwIs[vit-2::gfp]*), HAH52 (*pwIs23[vit-2::gfp]*), HAH53 (*adr-2(uu28);pwIs23[vit-2::gfp]*), HAH54 (*adr-2(G184R);pwIs23[vit-2::gfp]*), HAH55 (V5::SQD-1), HAH56 (V5::SQD-1;3xFLAG::ADR-2), HAH63 (*adr-1(tm668);pwIs23[vit-2::gfp]*).

The V5::SQD-1 strain (HAH55) was made using standard microinjection techniques and *dpy-10* co-CRISPR to segregate successful CRISPR via rolling F1 progeny and non-rolling F2 progeny (Ghanta and others 2021). Injection mix for the V5::SQD-1 strain (HAH55) included 1.5 µM Cas9 (IDT, Alt-R Cas9 nuclease V3), 4 µM tracrRNA (IDT), 3µM of crRNA (IDT) (HH3122) (**Table S1**), 1 μM *dpy-10* crRNA (IDT) (HH3118) (**Table S1**), 0.25 μM *dpy-10* SSODN (HH2448) (**Table S1**), and 0.75µM of repair template ssODN (HH3158) (**Table S1**) containing the V5::SQD-1 sequence. Expression of V5::SQD-1 was confirmed by immunoblotting.

Crossed strains were made by placing 10-15 males and 1 hermaphrodite on mating plates (NGM plates seeded with a small spot of *E. coli* OP50 in the center) and genotyping was performed for the F1 progeny and F2 progeny using primers mentioned in **Table S1**. The specific crosses performed included: creation of HAH51 by crossing HAH20 hermaphrodites to BB20 males, creation of HAH52 and HAH53 by crossing HAH20 hermaphrodites to *adr-2(uu28)* males, creation of HAH54 by crossing HAH51 hermaphrodites to HAH10 males, creation of HAH56 by crossing HAH58 hermaphrodites to HAH55 males, creation of HAH63 and HAH64 by crossing HAH20 hermaphrodites to BB21 males, creation of HAH65 by crossing BB20 males to HAH55 hermaphrodites.

### Brightfield and Fluorescence imaging

For **Figures 1A** and **3A**, imaged animals were mounted on 2% agarose pads in halocarbon oil and covered with a coverslip. Imaging was performed on a Zeiss SteREO Discovery V20. At least 10 animals were imaged per condition, representative images are shown in figures. For **Figures 5** and **6**, germlines were extruded from animals grown on RNAi as described below at 72 or 96 hours post egg-lay. Germlines were fixed with 1% paraformaldehyde and methanol and permeabilized with 0.2% Triton X-100 (Sigma-Aldrich). For **Figure 5B**, germlines were stained with SP56 primary antibody (a kind gift from Judith Kimble), followed by Alexa Fluor 594 (Thermo Fisher Scientific) and DAPI (4′,6-diamidino-2-phenylindole) (Thermo Fisher Scientific). For **Figure 5A** and **C** and **Figure 6**, germlines were stained with DAPI (4′,6-diamidino-2-phenylindole) (Thermo Fisher Scientific). Germlines were mounted in Vectashield (VectorLabs) and imaged using a Leica SP8 Scanning Confocal microscope (Indiana University Light Microscopy Imaging Center). Images were processed using ImageJ. For **Figure S5**, heterozygous null (GFP negative) animals of the balanced *sqd-1* null strain (VC1106) were selected from a synchronized population 96 hours post egg-lay. Germline extrusion, DAPI staining, and imaging were performed as described above. For **Figures 4 and 8**, germlines were extruded from synchronized L4 or day-one adult HH56 animals. Germlines were fixed with 1% paraformaldehyde and methanol and permeabilized with 0.2% Triton X-100 (Sigma-Aldrich). Germlines were stained with anti-V5 (Cell Signaling Technology) or anti-FLAG (Sigma-Aldrich) primary antibodies, followed by Alexa Fluor 594 (Thermo Fisher Scientific) or Alexa Fluor 488 (Thermo Fisher Scientific) and DAPI (4′,6-diamidino-2-phenylindole) (Thermo Fisher Scientific). For **Figure S7**, heterozygous null (GFP negative) animals of the balanced *sqd-1* null strain (VC1106) were selected from synchronized populations at 72 and 96 hours post egg-lay. Germlines were stained with DAPI and custom ADR-2 antibody (Rajendren and others 2018). For all confocal imaging experiments, germlines were mounted in Vectashield (VectorLabs) and imaged using a Leica SP8 Scanning Confocal microscope (Indiana University Light Microscopy Imaging Center). Images were processed using ImageJ.

**Figure 1:**
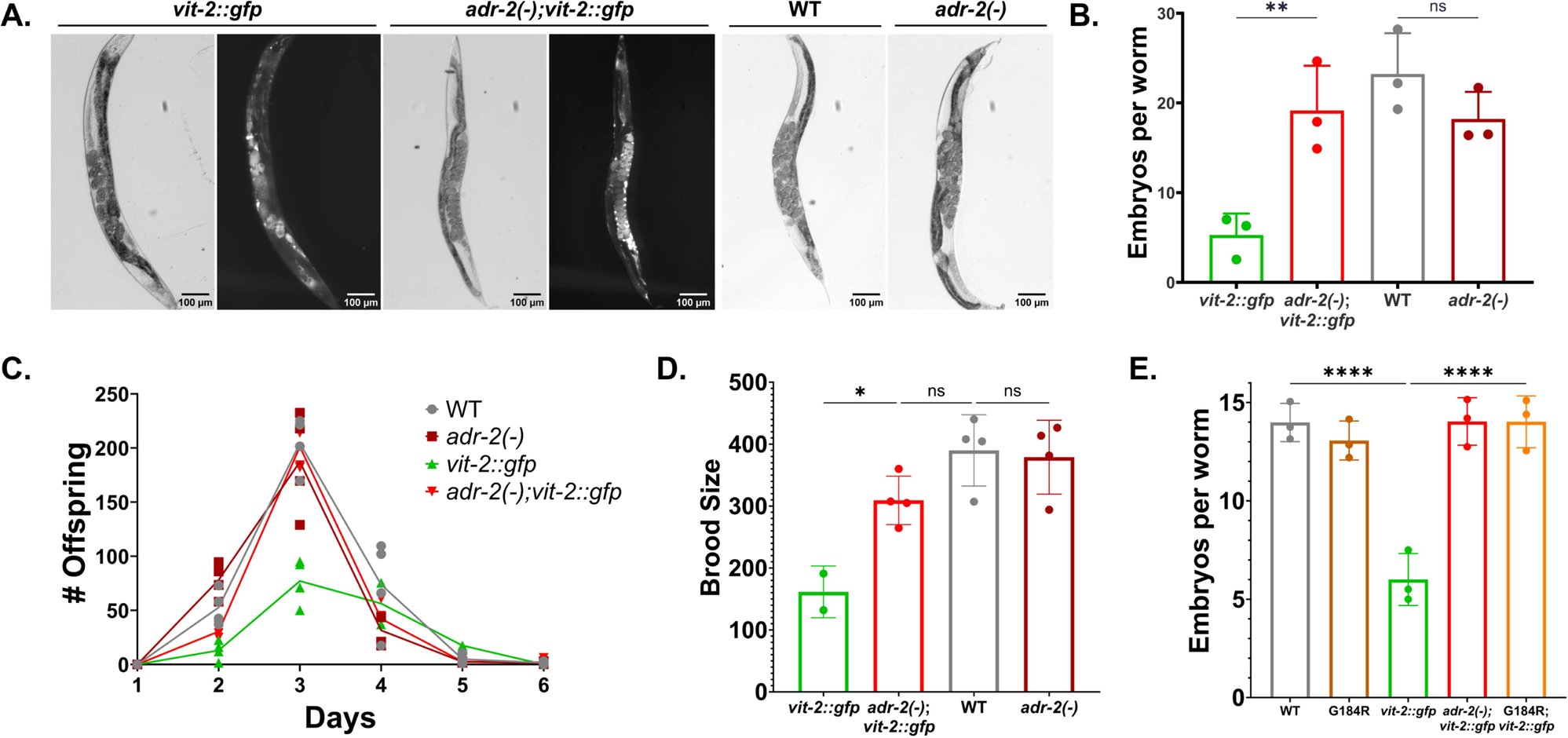
Loss of *adr-2* editing rescues the fertility defects of *vit-2::gfp* animals. A) Brightfield and fluorescence images in synchronized animals of the indicated genotypes. B) Embryo content of the indicated strains. Data represents 3 biological replicates, n=20 animals/replicate. Statistical significance determined via ordinary one-way Anova; ns= not significant (cut-off p≤0.05), ** indicates p≤0.01. C-D) Brood size assay of the indicated strains where both daily offspring (C) and total brood size (D) were plotted. Data represents 2-4 biological replicates. Statistical significance determined via ordinary one-way Anova; ns= not significant (p>0.05), * indicates p≤0.05. Connecting lines in panel C represent daily averages. E) Embryo content of the indicated strains. Data represents 3 biological replicates, n=20 worms/replicate. Statistical significance determined via ordinary one-way Anova; **** indicates p≤0.0001.

### Embryo content assay

Performed as previously described (Erdmann and others 2022). Briefly, animals were synchronized by plating 20 gravid adults on each plate and allowing them to lay eggs for 3 hours before removing. Synchronized offspring were allowed to develop to day 2 adulthood (except for the experiment in **Figure S1**, which was performed at day 1 adulthood). Twenty animals from each strain were placed in individual drops of 20% Arocep Ultra Bleach solution (Fisher), and the cuticles were allowed to dissolve for ∼10 minutes. The remaining embryos in each bleach drop were counted. The average embryo count from each set of 20 animals was calculated, representing one biological replicate. 3-5 biological replicates were performed for each strain.

### Brood size assay

Performed as previously described (Erdmann and others 2022). Animals were synchronized as stated above. Two L4-stage animals of each strain were plated on each of 10 NGM plates seeded with *E. coli* OP50. The animals were moved to fresh plates each day for the remainder of the experiment. Worms that died or crawled off the plate during the assay were excluded. Offspring laid on each plate were counted for two subsequent days after removal of the adults and divided by the number of adult animals plated on that plate, with adjustments made for animals that died or crawled off during the experiment. The experiment was ended when all adult animals produced no offspring for two consecutive days.

### Bleaching

Synchronized animals were obtained by bleaching gravid adult animals with a solution of 5M NaOH (33%) and Sodium Hypochlorite (Fisher Scientific) (66%). After the solution was added, animals were incubated on a shaker at 20°C for 7 minutes and then spun down to collect embryos. Embryos were washed with 1X M9 buffer (22 mM KH_2_PO_4_, 42.3 mM Na_2_HPO_4_, 85.6 mM NaCl, 1 mM MgSO_4_) solution thrice and incubated overnight in 1X M9 buffer at 20°C. The next day, hatched L1 worms were spun down and washed again with 1X M9 buffer thrice.

### RNA interference

Animals were synchronized and plated on RNAi plates (Plates with standard nematode growth medium, Ampicillin (50 μg/mL), Tetracycline (10 μg/mL), and IPTG (2mM)) seeded with HT115 bacteria containing RNAi vectors against various RNA binding proteins (RNAi vector collection a kind gift of Oliver Hobert, including plasmids compiled from the Ahringer and Vidal libraries) (**Table S2**). Each COPAS screening experiment also included animals fed with an empty RNAi vector and animals fed with an RNAi vector against GFP. After 3 days of growth at 20⁰C, the animals on each plate were washed off with 1X M9 buffer. The RNAi vector for each high-confidence regulator was confirmed by Sanger sequencing.

### GFP Quantification and data analysis

Animals were synchronized by bleaching and RNAi was performed as described above. Synchronized young adult animals were collected and run through the COPAS SELECT. The data were restricted to adult animals by eliminating all animals with time-of-flight values less than or equal to 120 or extinction values less than or equal to 176. The average GFP level for each sample was calculated, and values were normalized to the positive control (animals fed with an empty RNAi vector). RNAi vectors causing a >2.5 fold increase or decrease in GFP compared to the positive control were designated hits significantly affecting the phenotype. RNAi treatments that failed to produce young adult animals were excluded from screening.

### Quantitative Real-Time PCR

RNA interference was performed as described above. Young adult animals were washed off RNAi plates with 1X M9 buffer, pelleted, and frozen in Trizol. RNA was extracted using standard Trizol-Chloroform extraction followed by treatment with TURBO DNase (Ambion) and cleanup with the RNeasy Extraction Kit (Qiagen). Reverse transcription was performed on total RNA using random hexamer primers, oligo dT, and Superscript III (Thermo Fisher). Gene expression was determined using KAPA SYBR FAST Master Mix (Roche) and gene-specific primers (**Table S1**) on a Thermo Fisher Quantstudio 3 instrument. The primers designed for qPCR spanned an exon-exon junction to prevent detection of genomic DNA in the samples. For each gene analyzed, a standard curve of 8 to 10 samples of 10-fold serial dilutions of the amplified product were used to generate a standard curve of cycle threshold versus the relative concentration of amplified product. Standard curves were plotted on a logarithmic scale in relation to concentration and fit with a linear line. Fit (r^2^) values were around 0.99 and at least 7 data points fell within the standard curve. Each cDNA measurement was performed in 3 technical replicates, and each experiment was performed in 3 biological replicates.

### RNA sequencing library preparation

RNAi and RNA extraction from young adult animals was performed as described above. DNase-treated total RNA (4 μg) was subjected to two rounds of Poly-A selection using magnetic oligo(dT) beads (Invitrogen). Libraries were created from entire poly A-selected RNA samples using the KAPA RNA Hyperprep kit (Roche); cleanup steps were performed using AMPure XP beads (Beckman Coulter). Library fragment size distribution (200-900bp) was determined via Tapestation electrophoresis (Agilent). Libraries for each of three biological replicates per condition were pooled by concentration and sequenced on two NextSeq 2000 P2 (100 cycle) flow cells (total 600 million single-end reads) at the Indiana University Center for Genomics and Bioinformatics. The quality of the sequencing reads obtained was checked using FASTQC (version 0.12.1), the results of which are summarized in **Table S6**.

### Bioinformatic Analysis

Reads were aligned to a reference genome (*C. elegans* PRJNA13758 WS275) using STAR (version 2.7.10b). Percent uniquely mapped reads for each sample is listed in **Table S6**. Differential gene expression analysis was performed in R with DESeq2 (Love and others 2014). Full code for these analyses is available at: https://github.com/emierdma/GSF3587/tree/main. Representation factors in Figures 4 and 5 were calculated using the following formula: Rf = # of genes in common between groups A and B / ((# of genes in group A * # of genes in group B)/total # of genes in genome) (Kim and others 2001).

## Results

### Loss of A-to-I editing activity rescues the fertility defect of *vit-2::gfp* worms

A recent report has shown that dsRNA is imported into embryos along with yolk via receptor-mediated endocytosis (RME) (Marré and others 2016). As ADARs target dsRNA, we were interested in monitoring RME in relation to ADAR activity. To this aim, *adr* mutants were crossed into animals expressing a transgenic fusion of the vitellogenin gene *vit-2* with *gfp*, which were generated to monitor yolk uptake via RME (Grant and Hirsh 1999). Of note, we recently reported that this *vit-2::gfp* transgenic strain has a fertility defect resulting in fewer embryos in the uterus throughout reproductive adulthood and an overall reduced brood size (Erdmann and others 2022). Herein, we noted that when the *vit-2::gfp* transgene was crossed into an *adr-2* deletion mutant (*adr-2(ok735)*) (Hundley and others 2008), the animals appeared similar in size and population density to wild-type animals lacking the transgene. Brightfield and fluorescence imaging revealed that while *vit-2::gfp* animals contained roughly half the number of embryos as wild-type animals, *adr-2(−);vit-2::gfp* animals contained a similar number to wild-type and *adr-2(−)* animals lacking the *vit-2::gfp* transgene (**Figure 1A**). To quantify these potential differences, an embryo counting assay was performed. The average number of embryos contained in the uterus of *adr-2(−);vit-2::gfp* animals was significantly greater than that of *vit-2::gfp* animals (**Figure 1B**). However, the effect of *adr-2* on embryo content is specific to the *vit-2::gfp* transgenic background, as *adr-2(−)* animals did not show a significant difference in embryo content compared to wild-type animals (**Figure 1B**). To ensure that the restoration of fertility was not due to any background mutations in the *adr-2(ok735)* strain or the specific *adr-2* genetic lesion, the *vit-2::gfp* transgene was crossed into a second *adr-2* deletion mutant (*adr-2(uu28)* (Reich and others 2018)). A second deletion allele was used as attempts to create a transgenic rescue for *adr-2* have been unsuccessful to date, likely due to its native location in a six-gene operon. Similar to *adr-2(ok735)*, *adr-2(uu28);vit-2::gfp* animals had significantly greater embryo content than *vit-2::gfp* animals (**Figure S1**), and the *adr-2(uu28)* animals without the *vit-2::gfp* transgene did not significantly differ from wild-type animals. These results indicate that loss of *adr-2* is sufficient to rescue the embryo content of animals expressing the *vit-2::gfp* transgene.

To gain further insight into how loss of *adr-2* promoted fertility in the *vit-2::gfp* animals, we tested whether the differences observed in the single time-point embryo content resulted from differences in reproductive timing. To test this, the number of viable offspring produced by each strain during each day of reproductive adulthood was monitored. Egg-laying for all strains started, peaked, and ended on the same days, suggesting no significant changes in reproductive timing in adulthood (**Figure 1C**). With respect to total brood size over the six peak days of egg-laying, *adr-2(−);vit-2::gfp* animals had a significantly greater average brood size compared to *vit-2::gfp* animals, and had a similar average brood size to non-transgenic strains (**Figure 1D**). Furthermore, there was no significant difference between the average brood sizes of wild-type and *adr-2(−)* animals, confirming that this effect of *adr-2* on fertility is transgene-specific. Taken together, these results indicate that the loss of *adr-2* is sufficient to rescue the fertility defects of animals expressing the *vit-2::gfp* transgene.

To begin to investigate how loss of *adr-2* relieves the fertility defect in *vit-2::gfp* animals, we tested whether loss of A-to-I RNA editing is sufficient to rescue embryo content. The *vit-2::gfp* transgene was crossed into a strain expressing ADR-2 containing a point mutation (G184R) that abolishes deamination activity while retaining RNA binding capabilities (Deffit and others 2017). Similarly to *adr-2(−);vit-2::gfp* animals, *adr-2(G184R);vit-2::gfp* animals show similar embryo counts to non-transgenic animals (**Figure 1E**), suggesting that loss of editing by ADR-2 is sufficient to rescue fertility in *vit-2::gfp* animals.

Editing by ADR-2 can be regulated by the deaminase-deficient cofactor ADR-1, the other ADAR family member in *C. elegans* (Arribere and others 2020). Because the fertility phenotype is editing-dependent, we sought to determine whether *adr-1* affects fertility in *vit-2::gfp* animals, and an *adr-1(−);vit-2::gfp* strain was created. In contrast with *adr-2*, loss of *adr-1* alone did not restore fertility in *vit-2::gfp* animals (**Figure S2**). These data are consistent with the observation that the rescue of fertility observed upon loss of *adr-2* is not simply due to loss of RNA binding by any ADAR family member. In addition, these data also provide additional evidence that the enhanced fertility of the *vit-2::gfp* strain observed upon crossing with *adr-2* deletions is not due to removal of an additional homozygous mutation in these genetic strains. Together, our data suggest a model wherein loss of RNA editing by ADR-2 improves fertility of *vit-2::gfp* animals independent of ADR-1.

### Screening for *adr-2*-related embryogenesis and RME defects

While the above data alone may reflect a transgenic background-specific effect of *adr-2* on fertility, alongside previous reports of synthetic fertility phenotypes for ADAR mutant animals (Fischer and Ruvkun 2020; Reich and others 2018), it could also suggest a role for *adr-2* in regulating embryogenesis. However, like many factors regulating reproduction (Vanden Broek and others 2022), loss of *adr-2* alone does not cause changes in fertility in wild-type animals (**Figure 1**). To gain a better understanding of how *adr-2* may interact with embryogenic and/or RME pathways, we designed a screen to identify factors that impact embryogenesis and RME in *adr-2(−)* animals. Considering that proximal oocytes and embryos in *vit-2::gfp* animals are fluorescent, and that the fertility defect results in fewer embryos contained within the uterus, we reasoned that fertility in *vit-2::gfp* animals could be estimated by measuring the fluorescence intensity of gravid worms. The COPAS SELECT (Union Biometrica) is a large particle sorter that can measure the length, density, and fluorescence intensity of large populations of individual animals. Sorting with COPAS instruments has been used previously to measure the impact of various chemicals on embryos within adult *C. elegans* (Shin and others 2019). However, it should also be noted that as previous studies have shown that disruptions in vitellogenin provisioning can cause an accumulation of yolk in the pseudocoelom of the animal (Grant and Hirsh 1999), COPAS sorting could also detect higher GFP signals that occur due to defective RME.

To test this screening method, we used COPAS sorting to measure the fluorescence intensity of large populations of gravid *vit-2::gfp* and *adr-2(−);vit-2::gfp* animals. Synchronized *vit-2::gfp* animals and *adr-2(−);vit-2 gfp* animals were allowed to develop for 60 hours prior to sorting. The length and density measures were used to restrict the analysis to adult animals, and the fluorescence values were averaged for each sorting run, which ranged from ∼2000-5000 animals. The average fluorescence intensity of adult *adr-2(−);vit-2::gfp* animals was significantly higher than that of *vit-2::gfp* animals (**Figure 2A**), mirroring the embryo content and brood size of the strains (**Figure 1**).

**Figure 2:**
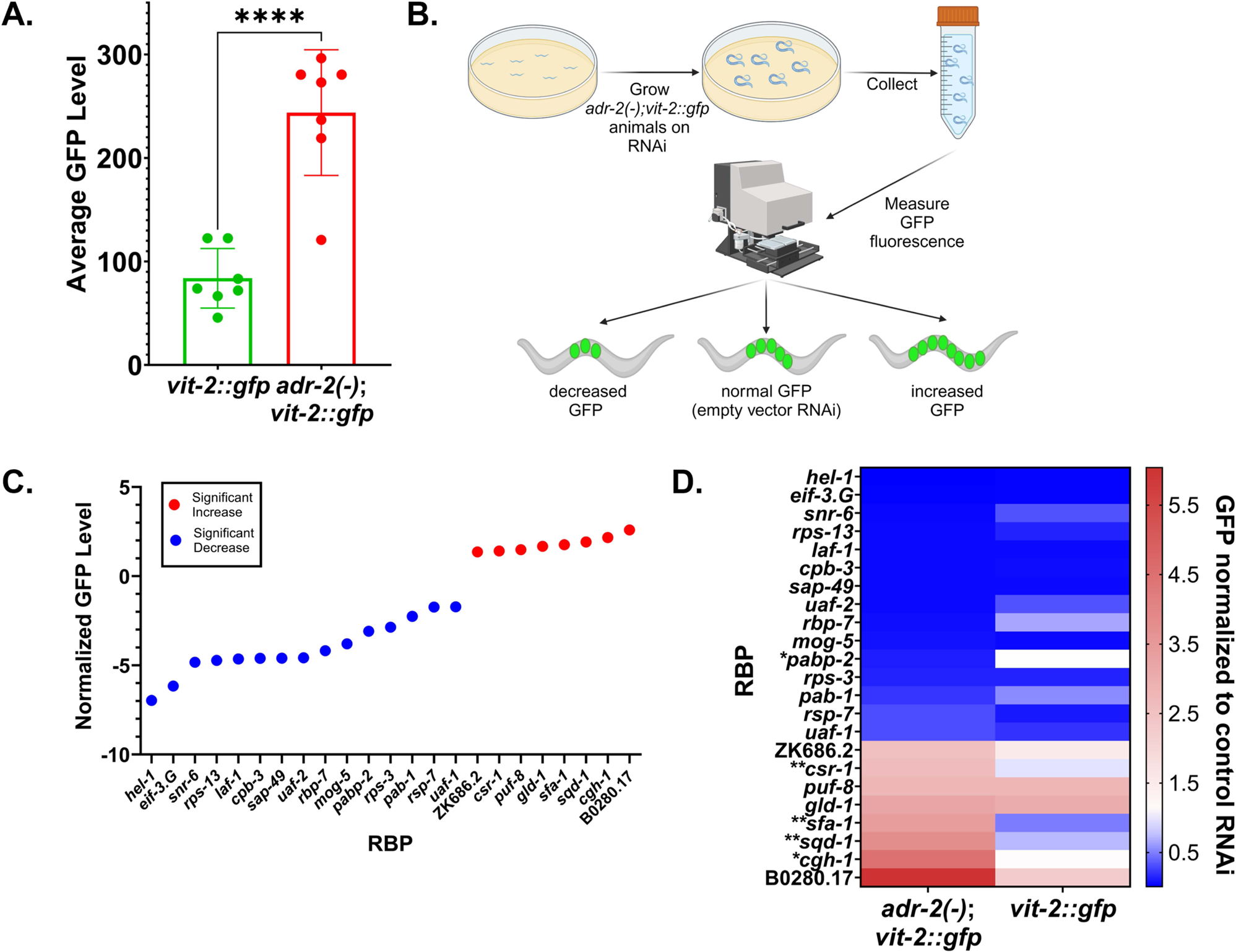
Screen for RNA binding proteins that impact vitellogenesis and embryo production. A) COPAS fluorescence quantification of synchronized *vit-2::gfp* and *adr-2(−);vit-2::gfp* worms. Data represents 7 biological replicates, ∼2000-5000 worms per replicate. Statistical significance determined via unpaired t-test; **** indicates p≤0.0001. B) Schematic of screen. Figure created with BioRender. C) High confidence modulators of GFP level in *adr-2(−);vit-2::gfp* animals. Based on results in **Figure S3A-B**. D) Heat map showing fluorescence changes upon RNAi of each high-confidence modulator in *adr-2(−);vit-2::gfp* animals compared to *vit-2::gfp* animals. * indicates RNAi treatments causing a significant change in GFP compared to controls in *adr-2(−);vit-2::gfp* animals but no significant change from controls in *vit-2::gfp* animals. ** indicates RNAi treatments causing a significant change in GFP compared to controls in *adr-2(−);vit-2::gfp* animals and a change in the opposite direction in *vit-2::gfp* animals.

As previous synthetic embryogenesis defects have been shown when ADARs are lost along with other RNA binding proteins (RBPs) (Reich and others 2018) (Fischer and Ruvkun 2020), we decided to focus our screen on RBPs. In a primary screen to identify factors affecting embryo content and/or vitellogenin provisioning, *adr-2(−);vit-2::gfp* animals were synchronized and treated with RNA interference (RNAi) against 255 RBPs that represent diverse classes of RBPs, including those harboring domains that recognize single stranded RNA, such as the K Homology (KH) and RNA Recognition Motif (RRM) domains as well as those that recognize double-stranded RNA (dsRNA), such as the dsRNA binding domain (dsRBD) (**Table S2**). After 60 hours, animals were collected from each treatment and the average GFP fluorescence of each strain was measured using the COPAS SELECT, restricting the analysis to adult animals as specified above (**Figure 2B**). Each experiment included a control group fed with bacteria expressing an empty RNAi vector and a control group treated with *gfp* RNAi. Average fluorescence values for each experimental treatment of ∼2000-5000 animals were normalized to the average fluorescence value for the *adr-2(−);vit-2::gfp* animals treated with empty vector RNAi. Treatments that caused a 2.5-fold or greater change in fluorescence were selected as hits (**Figure S3A, Table S3**); however, it should be noted that factors not meeting this threshold should not be ruled out as potential regulators of fertility and/or vitellogenin provisioning, as variable efficacy of RNAi vectors may preclude some factors from meeting the screening threshold. Each hit was tested in a second, independent biological replicate (**Figure S3B, Table S4**), and those that did not show a similar trend in both replicates were eliminated to allow for identification of only high-confidence regulators of embryo content or vitellogenin provisioning phenotypes (**Figure 2C**). Of the 255 RBPs tested, 23 high-confidence regulators were identified (**Figure 2C**). Importantly, several known regulators of fertility or RME were captured in the screen (**Table S5**), including *gld-1* (Lee and Schedl 2010) and *puf-8* (Datla and others 2014).

To narrow our search to *adr-2*-dependent phenotypes, *vit-2::gfp* animals were also subjected to RNAi against the high-confidence regulators (**Table S4**). It is important to note that because the *vit-2::gfp* strain has low baseline fertility, it is unlikely to identify changes in GFP fluorescence as drastic as those seen in *adr-2(−);vit-2::gfp* animals, especially for factors that decrease fluorescence. Thus, we compared the general trend of fluorescence in *vit-2::gfp* animals to *adr-2(−);vit-2::gfp* animals (**Figure 2D**). For the majority of the RBPs (18/23), when compared to the control RNAi, a similar change in GFP fluorescence was observed in both *vit-2::gfp* and *adr-2(−);vit-2::gfp* animals, suggesting the changes in fluorescence are independent of *adr-2*. The 5 other RBPs fell into one of two categories. The first category (denoted with * in **Figure 2D**) includes those that caused a significant change in GFP fluorescence in *adr-2(−);vit-2::gfp* animals but produced almost no difference in GFP fluorescence in *vit-2::gfp* animals (*pabp-2* and *cgh-1*). The second category (denoted with ** in **Figure 2D**) includes those that had an opposite effect on GFP fluorescence in *vit-2::gfp* animals compared to *adr-2(−);vit-2::gfp* animals (*sfa-1*, *csr-1*, and *sqd-1*) (**Figure 2D**). Because *pabp-2* RNAi caused decreased fluorescence in *adr-2(−);vit-2::gfp* animals, the lack of a significant change in GFP fluorescence in the *vit-2::gfp* animals may be attributable to the already lowered fertility in this strain. However for *cgh-1* RNAi, which causes an increase in fluorescence in *adr-2(−);vit-2::gfp* animals, the lack of a similar increase in *vit-2::gfp* animals suggests a synthetic interaction between *cgh-1* and *adr-2*. Similarly, RNAi treatments for *sfa-1*, *csr-1*, and *sqd-1* showed a significant increase in GFP fluorescence in *adr-2(−)::vit-2::gfp* and a slight decrease in fluorescence in *vit-2::gfp* animals, suggesting potential synthetic interactions of *sfa-1*, *csr-1* and *sqd-1* with *adr-2*. Together, this screen identified 23 high-confidence regulators of fertility and/or yolk provisioning in *C. elegans*, as well as four RBPs that may have synthetic interactions with ADR-2.

### Investigating potential synthetic interactions with *adr-2*

To further investigate the potential synthetic interactions of *adr-2* with *cgh-1*, *csr-1*, *sfa-1*, and *sqd-1*, we first sought to determine whether the *adr-2*-dependent changes in GFP fluorescence observed in the screen resulted from different numbers of embryos or from changes in yolk provisioning. To investigate this, imaging was performed on *vit-2::gfp* and *adr-2(−);vit-2::gfp* worms treated with RNAi against each factor (**Figure 3A**). Consistent with the embryo counting results (**Figure 1B**), the brightfield images revealed that *vit-2::gfp* animals contain fewer embryos than *adr-2(−);vit-2::gfp* animals (top brightfield panels, **Figure 3A**). Focusing on the *vit-2::gfp* animals, those treated with RNAi for *cgh-1*, *csr-1*, *sfa-1* or *sqd-1* all contained fewer embryos than *vit-2::gfp* animals treated with control RNAi (**Figure 3A**), consistent with previous reports of fertility defects observed upon loss of *cgh-1* (Audhya and others 2005), *csr-1* (Yigit and others 2006) and *sqd-1* (Maeda and others 2001). In keeping with these fertility defects, *adr-2(−);vit-2::gfp* animals treated with RNAi against *cgh-1*, *csr-1*, and *sqd-1* appeared to contain fewer embryos than *adr-2(−);vit-2::gfp* animals treated with control RNAi. Of note, for all three of these factors, the *adr-2(−);vit-2::gfp* animals treated with RNAi contained more embryos than *vit-2::gfp* animals treated with RNAi, suggesting a possible source of the increased GFP intensity observed in the COPAS screen and possible *adr-2*-dependent effects of *cgh-1*, *csr-1* and *sqd-1* on fertility. In contrast, *adr-2(−);vit-2::gfp* animals treated with *sfa-1* RNAi had a similar embryo content to *adr-2(−);vit-2::gfp* animals treated with control RNAi (**Figure 3A**), suggesting that *sfa-1* does not affect fertility in this genetic background.

**Figure 3:**
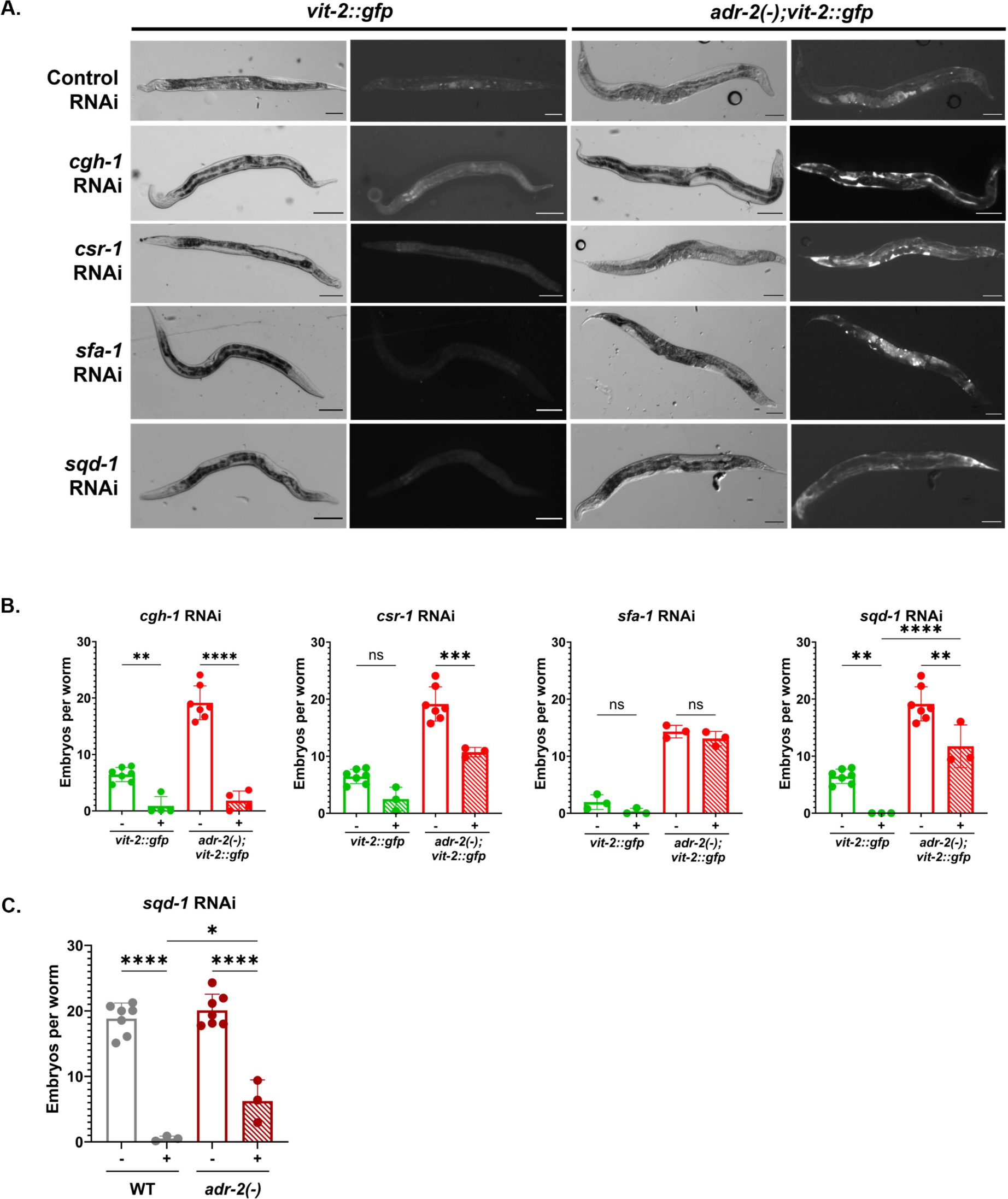
RNAi against *cgh-1*, *csr-1*, *sfa-1*, and *sqd-1* causes accumulation of vitellogenin in the body cavity of *adr-2(−);vit-2::gfp* animals A) Representative brightfield and fluorescent images of *vit-2::gfp* and *adr-2(−);vit-2::gfp* animals treated with control RNAi and RNAi against *cgh-1*, *sfa-1*, *sqd-1*, and *csr-1*. Scale bars = 100 μm. B) Embryo content of *vit-2::gfp* and *adr-2(−);vit-2::gfp* animals treated with empty vector RNAi or RNAi against *csr-1*, *sfa-1*, and *sqd-1*. Synchronized adult animals were placed individually in 20% bleach until the cuticles dissolved, and the remaining embryos from each worm were counted. Data represents 3 biological replicates, 20 animals/replicate. Significance determined via One-Way ANOVA; ns = not significant (p>0.05), ** indicates p≤0.01, *** indicates p≤0.001, **** indicates p≤0.0001. C) Embryo content of wild-type and *adr-2(−)* animals treated with empty vector RNAi or *sqd-1* RNAi. Synchronized adult animals were placed individually in 20% bleach until the cuticles dissolved, and the remaining embryos from each worm were counted. Data represents 3 biological replicates, 20 animals/replicate. Significance determined via One-Way ANOVA; * indicates p≤0.05, **** indicates p≤0.0001.

When focusing on the fluorescence images, for *vit-2::gfp* animals treated with RNAi against *cgh-1*, *csr-1*, *sfa-1*, and *sqd-1*, aside from the reduced oocytes, there does not appear to be a major change in GFP fluorescence intensity or localization compared to *vit-2::gfp* animals treated with control RNAi (**Figure 3A**, left panels). These data are consistent with the GFP fluorescence intensity observed using the COPAS instrument (**Figure 2D**, right panel). For the *adr-2(−);vit-2::gfp* animals treated with control RNAi, an increased number of GFP positive embryos is observed in the fluorescence images when compared to *vit-2::gfp* animals under the same treatment (**Figure 3A**, top panels). These data are consistent with the COPAS GFP intensity data (**Figure 2A**) and the images and embryo counts performed on these strains in the absence of RNAi (**Figure 1A, B**). Strikingly, for the *adr-2(−);vit-2::gfp* animals treated with RNAi against *cgh-1, csr-1, sfa-1,* or *sqd-1*, the GFP fluorescence was not restricted to the embryos and was observed throughout the body cavity (**Figure 3A**, right panels). This may at least partially account for the observed increase in GFP fluorescence intensity using the COPAS instrument for *adr-2(−);vit-2::gfp* animals treated with RNAi against each factor (**Figure 2C**). Of note, alterations in yolk protein localization in animals treated with RNAi to *sfa-1* and *cgh-1* have been previously reported (Balklava and others 2007). Because the accumulation of VIT-2::GFP fluorescence in the body cavity of *adr-2(−);vit-2::gfp* animals treated with RNAi against *cgh-1*, *csr-1*, *sfa-1*, and *sqd-1* is not seen in *vit-2::gfp* animals with these same treatments, these observations suggest that *adr-2* affects vitellogenin provisioning in these animals, either by altering uptake of yolk into oocytes via RME or by altering vitellogenin expression levels. However, it is possible that yolk provisioning is still disrupted in *vit-2::gfp* animals treated with *cgh-1*, *csr-1*, *sfa-1*, and *sqd-1*, but these defects are not apparent due to the low baseline expression of VIT-2::GFP in these animals compared to *adr-2(−);vit-2::gfp* animals.

To more quantitatively look at fertility in these backgrounds, embryo content assays were performed for *vit-2::gfp* and *adr-2(−);vit-2::gfp* animals treated with RNAi for each RBP analyzed above. For both *vit-2::gfp* and *adr-2(−);vit-2::gfp* animals, treatment with *cgh-1* RNAi caused a significant decrease in embryo content (**Figure 3B**, left panel). This suggests that, while treatment with *cgh-1* RNAi results in altered vitellogenin accumulation/expression in the absence of *adr-2* (**Figure 3A**), CGH-1 affects fertility independent of *adr-2* (**Figure 3B**, left panel). Consistent with the brightfield images, treatment with *csr-1* RNAi caused a decrease in embryo content in both strains, but reached statistical significance only in the *adr-2(−);vit-2::gfp* animals (**Figure 3B**, left middle panel). These data suggest that, like *cgh-1*, *csr-1* RNAi results in altered vitellogenin accumulation/expression in the absence of *adr-2* but reduces fertility independent of *adr-2*.

Treatment of *vit-2::gfp* animals with *sfa-1* RNAi did not cause a statistically significant change in embryo content (**Figure 3B**, right middle panel); however, given the already low embryo content of *vit-2::gfp* animals, even the slight reduction might account for the lack of embryos observed in the images of *vit-2::gfp* animals treated with *sfa-1* RNAi (**Figure 3A**). In *adr-2(−);vit-2::gfp* animals, *sfa-1* RNAi treatment did not significantly alter embryo content (**Figure 3B**, right middle panel), consistent with the imaging data. In sum, these data suggest that *sfa-1* does not significantly impact fertility, and that the increased GFP fluorescence observed upon *sfa-1* RNAi treatment reflects a change in accumulation/expression of VIT-2::GFP within the body cavity as observed, or additionally in the oocytes and embryos.

Consistent with previous reports of sterility in animals lacking *sqd-1* (Maeda and others 2001), treatment of *vit-2::gfp* animals with *sqd-1* RNAi resulted in an absence of embryos (**Figure 3B**, right panel). The *adr-2(−);vit-2::gfp* animals treated with *sqd-1* RNAi exhibited a reduced embryo content compared to control, but interestingly, these animals were no longer sterile. In fact, *adr-2(−);vit-2::gfp* animals treated with *sqd-1* RNAi contained significantly more embryos than *vit-2::gfp* animals treated with *sqd-1* RNAi (**Figure 3B**, right panel). To more broadly assess whether loss of *adr-2* restores fertility to animals depleted for *sqd-1*, embryo content assays were performed on non-transgenic, wild-type and *adr-2(−)* animals treated with control or *sqd-1* RNAi. In wild-type animals, we see the previously reported sterility caused by loss of *sqd-1* (**Figure 3C**). In *adr-2(−)* animals, *sqd-1* RNAi also causes a reduction in embryo content. However, *sqd-1* RNAi does not result in complete sterility in *adr-2(−)* animals, and similar to the observations in the transgenic background, *adr-2(−)* animals treated with *sqd-1* RNAi show a significant increase in embryo content compared to wild-type animals treated with *sqd-1* RNAi (**Figure 3C**). These results are intriguing, as they suggest that *adr-2* may contribute to the fertility defects observed in wild-type animals depleted for *sqd-1*. However, it is also possible that fertility is enhanced in the *adr-2(−)* background due to reduced efficiency of RNAi, leading to a higher expression of *sqd-1* mRNA in *adr-2(−)* animals treated with *sqd-1* RNAi. To rule out this possibility, quantitative real-time PCR was performed to measure *sqd-1* levels in total RNA isolated from wild-type and *adr-2(−)* animals treated with control or *sqd-1* RNAi. Importantly, both strains showed similar reduced levels of *sqd-1* mRNA when treated with *sqd-1* RNAi (**Figure S4**). Thus, these data reflect a role for *adr-2* in modulating fertility in animals depleted of *sqd-1*.

### Loss of sqd-1 disrupts the switch from spermatogenesis to oogenesis in young adult animals

To investigate how *adr-2* may modulate fertility in *sqd-1* depleted animals, we first needed to understand the root of the fertility defects caused by *sqd-1* RNAi treatment. As SQD-1 function has not been studied in *C. elegans*, we first sought to determine whether SQD-1 was expressed in the germline. To assess SQD-1 localization, we installed a V5 epitope-tag at the N-terminus of SQD-1 using CRISPR genome engineering. Next, dissected germlines were stained for SQD-1 and DAPI and imaged. SQD-1 staining is observed throughout the undifferentiated germ cells and oocytes in oogenic young adult germlines (**Figure 4A**). SQD-1 staining appears in the cytoplasm of these cells and is largely absent from the nuclei, where DAPI staining is present (**Figure 4A**). SQD-1 staining is absent in the spermatheca in young adult animals (**Figure 4A**), suggesting SQD-1 is not expressed in sperm, which are distinguishable as small, highly condensed DAPI foci in the proximal gonad. To understand where SQD-1 is expressed in spermatogenic germlines, SQD-1 staining was also performed in dissected germlines from L4 stage animals. Similar to young adults, SQD-1 is expressed in the cytoplasm of undifferentiated germ cells during spermatogenesis, but is not observed in sperm (**Figure 4B**). To our knowledge, this is the first demonstration of SQD-1 expression in the germline of *C. elegans*.

**Figure 4:**
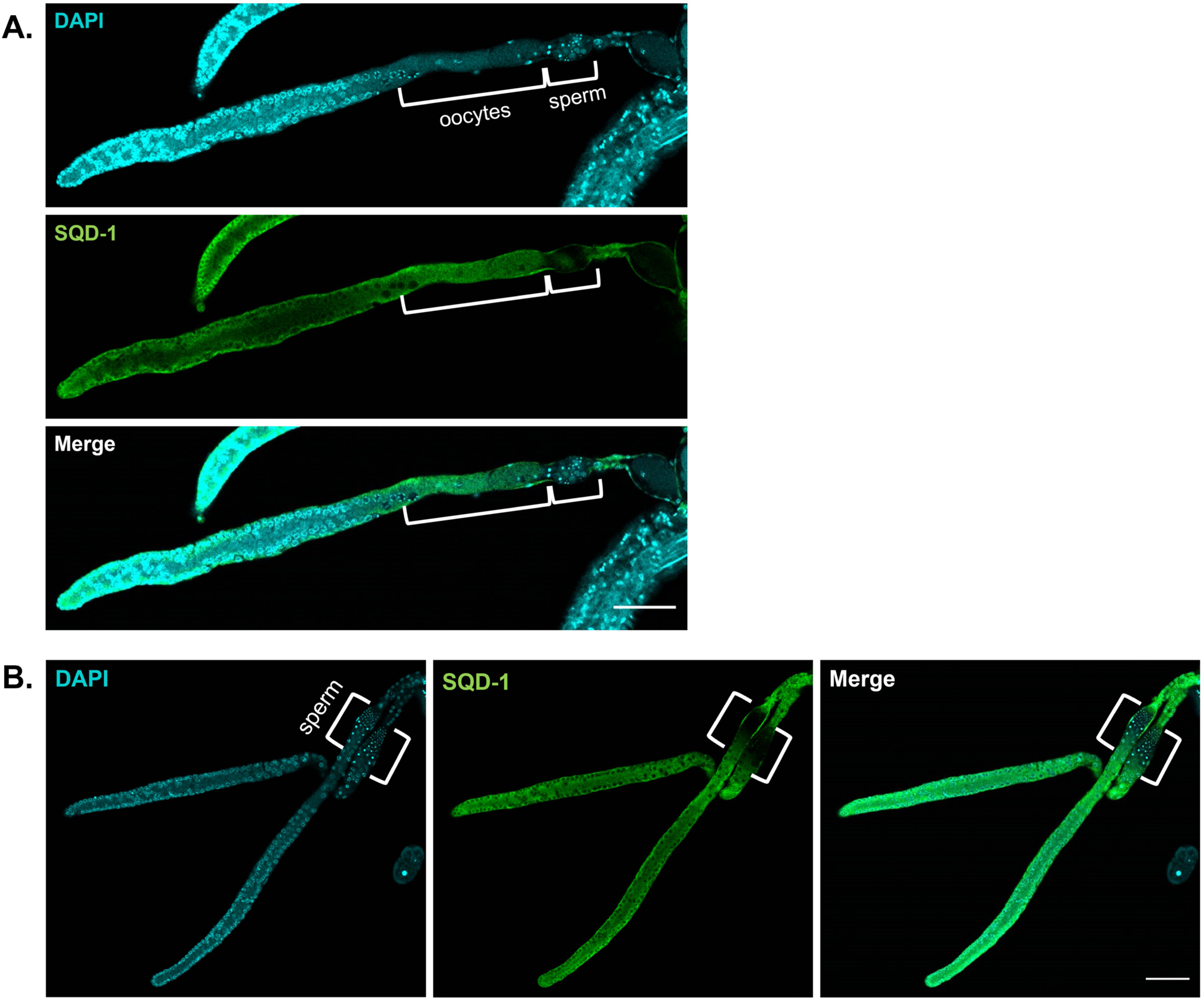
SQD-1 germline expression patterns. A) Representative confocal images of dissected germlines from young adult (oogenic) animals stained for DAPI (cyan) and SQD-1 (green). Scale bars = 50 μm. Brackets denote oocytes and sperm. B) Representative confocal images of dissected germlines from L4 (spermatogenic) animals stained for DAPI (cyan) and SQD-1 (green). Scale bars = 50 μm. Brackets denote sperm.

Next, we assessed the effects of depletion of *sqd-1* on germline morphology. Germline dissection and nuclear staining with DAPI were performed on wild-type animals treated with control and *sqd-1* RNAi. At 72 hours post egg-lay, wild-type animals treated with control RNAi produced typical germline structures (Pazdernik and Schedl 2013), with many syncytial nuclei at the distal end of the germline, apparent germ cell culling around the bend of the gonad, and individual, cellularized oocytes in the proximal germline region (**Figure 5A**, top panel, **Figure 5D**). Wild-type, day-one adults (72 hrs post egg-lay) treated with *sqd-1* RNAi produced germlines that appeared normal at the distal end with many syncytial nuclei, but nearly 90% of germlines did not form individual oocytes at the proximal end. Instead, the proximal ends were populated by condensed nuclei, resembling spermatogenic germlines (**Figure 5A**, bottom panel, **Figure 5D**). Staining with sperm marker SP56 confirmed that the proximal gonads of the *sqd-1* depleted animals are populated by sperm, while SP56 staining was not seen in the proximal gonad of animals treated with control RNAi (**Figure 5B**). On day two of adulthood (96 hrs post egg-lay), wild-type animals treated with *sqd-1* RNAi presented a range of germline defects. About one-third showed no obvious germline defect, resembling wild-type animals treated with control RNAi with oocytes present in the proximal gonad and sperm constrained to the spermatheca (**Figure 5C**, top panel, **Figure 5D**). Intermediate defects were seen in ∼30% of germlines, with oocytes present in the mid-proximal gonad but an overpopulation of sperm extending out of the spermatheca and into the proximal gonad (**Figure 5C**, middle panel, **Figure 5D**). Finally, ∼40% showed severe defects, with no distinguishable oocytes and proximal gonads populated by sperm (**Figure 5C**, bottom panel, **Figure 5D**). Together, these data suggest that loss of *sqd-1* results in a failure to switch from spermatogenesis to oogenesis in young adult animals, leading to an accumulation of sperm and infertility. The range of severity of defects at the later timepoint may reflect a decrease in the efficacy of *sqd-1* RNAi in older animals, allowing some animals to reach a threshold of *sqd-1* expression that allows oogenesis. Alternatively, it is possible that loss of *sqd-1* delays but does not abolish the switch from spermatogenesis to oogenesis. However, as ∼40% of animals treated with *sqd-1* RNAi still exhibit a failure to switch to oogenesis at the later timepoint, it is likely the differences in phenotype reflect differences in RNAi penetrance rather than timing defects. To directly address this issue, *sqd-1* null germlines derived from a balanced deletion mutant strain were imaged at this same timepoint (96 hours post egg-lay). All *sqd-1* null germlines imaged showed a severe defect, with the proximal germline entirely populated by sperm and no oocytes present (**Figure S5**). These observations confirm that the defects caused by loss of *sqd-1* are consistent with a failure to switch from spermatogenesis to oogenesis, rather than a delayed switch. Together with our localization data, these data suggests that SQD-1 expression may be necessary to specify oocyte fate or suppress sperm fate in differentiating germ cells.

**Figure 5:**
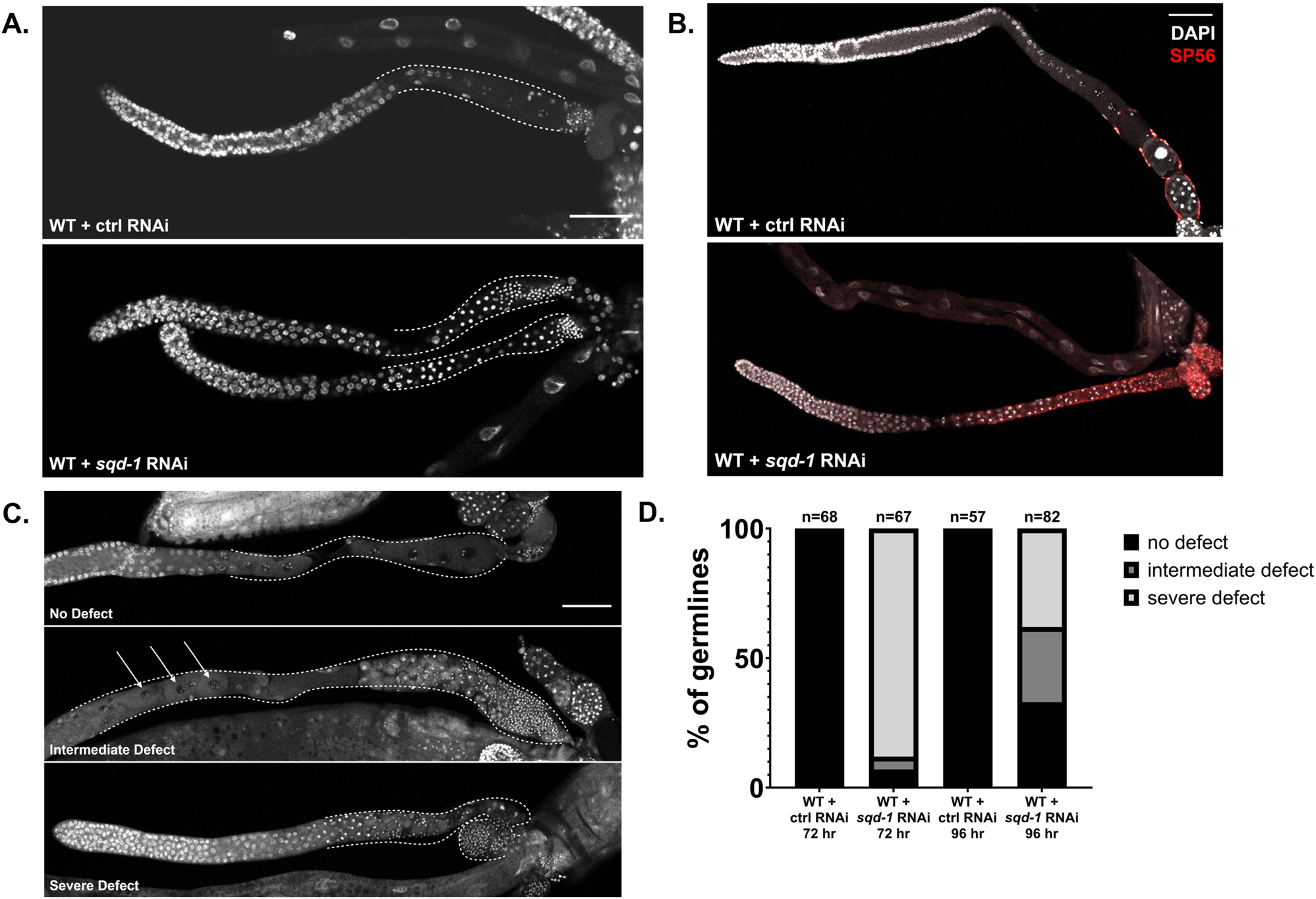
Loss of *sqd-1* causes defects in the developmental switch from spermatogenesis to oogenesis. A) Representative confocal images of dissected germlines from wild-type, 72 hr post egg-lay animals treated with control RNAi or *sqd-1* RNAi, stained with DAPI. Scale bar = 50 μm. Dotted lines indicate proximal germline region. B) Representative confocal images of dissected germlines from wild-type, 72 hr post egg-lay animals treated with control RNAi or *sqd-1* RNAi. Stained with DAPI (gray) and sperm marker SP56 (red). Scale bar = 50 μm. C) Representative examples of no defect, intermediate defect, and severe defect in wild-type, 96 hr post egg-lay animals treated with RNAi against *sqd-1*. Stained with DAPI. Scale bar = 50 μm. Dotted lines indicate proximal germline region. Arrows indicate location of oocytes in intermediate defect. D) Distribution of germline defects in wild-type animals treated with control or *sqd-1* RNAi, 72 hr or 96 hr post egg-lay. n for each condition is indicated above bars.

### The loss of adr-2, but not RNA editing, partially restores oogenesis in animals lacking sqd-1

Next, we aimed to address the role of *adr-2* in modulating fertility in *sqd-1* depleted animals. Due to the partial rescue of embryo numbers in the *adr-2(−)* animals treated with *sqd-1* RNAi (**Figure 3C**), we expected that the germline defects caused by *sqd-1* RNAi would be less severe in *adr-2(−)* animals than wild-type animals. Thus, we assessed germline morphology in this strain. When *sqd-1* RNAi is performed in *adr-2(−)* animals, the germlines do not show the severe defects observed in wild-type animals treated with *sqd-1* RNAi. At 72 hours post egg-lay, most *adr-2(−)* germlines treated with *sqd-1* RNAi (82%) appear to proceed through oocyte and embryo production as normal, with no obvious morphological differences from wild-type and *adr-2(−)* germlines treated with control RNAi (**Figure 6A,B**). In contrast to wild-type animals treated with *sqd-1* RNAi, only small portion of *adr-2(−)* germlines treated with *sqd-1* RNAi showed either intermediate defects (sperm and oocytes in the proximal gonad) (8%) or severe defects (only sperm in the proximal gonad) (10%) (**Figure 6B**). These data suggest that loss of *adr-2* can rescue the switch from spermatogenesis to oogenesis in *sqd-1* depleted animals. However, as we previously observed decreased embryo content in *adr-2(−)* animals treated with *sqd-1* RNAi compared to controls, it seems that this rescue of oogenesis is not sufficient to fully restore fertility.

**Figure 6:**
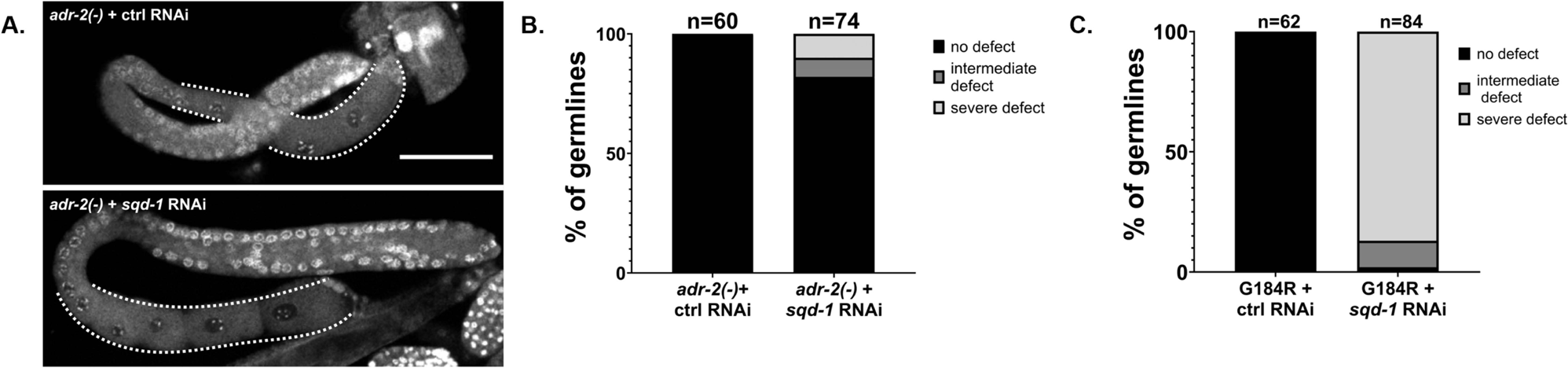
Loss of *adr-2* restores oogenesis in *sqd-1* depleted animals. A) Representative confocal image of dissected germlines from *adr-2(−),* 72 hr post egg-lay animals treated with control or *sqd-1* RNAi, stained with DAPI. Scale bar = 50 μm. Dotted lines indicate proximal germline region. B) Distribution of germline defects in *adr-2(−)* animals treated with control or *sqd-1* RNAi, 72 hr post egg-lay. n for each condition is indicated above bars. C) Distribution of germline defects in *adr-2(G184R)* animals treated with control or *sqd-1* RNAi, 72 hr post egg-lay. n for each condition is indicated above bars.

To begin to understand how ADR-2 may be impacting fertility in animals depleted for *sqd-1*, similar to our analysis of *vit-2::gfp*, we tested whether the *sqd-1* RNAi germline defects are dependent on the RNA editing function of ADR-2. Animals expressing the editing-deficient *adr-2* mutant (G184R) (Deffit and others 2017) were fed with either control or *sqd-1* RNAi. All *adr-2(G184R)* animals fed with control RNAi displayed normal oogenic germline morphology at 72 hours post egg-lay, but *adr-2(G184R)* animals fed with *sqd-1* RNAi displayed similar levels (87%) of spermatogenic germlines (**Figure 6C**) as wild-type animals fed with *sqd-1* RNAi (**Figure 5D**). Intermediate defects were seen in 11% of *adr-2(G184R)* germlines treated with *sqd-1* RNAi, and 2% displayed wild-type morphology (**Figure 6C**). Because loss of editing does not rescue *sqd-1* germline defects as loss of *adr-2* does, this suggests that the defects are independent of RNA editing by ADR-2, and likely depend on RNA binding by ADR-2.

### Loss of *adr-2* partially restores oogenic gene expression in animals lacking *sqd-1*

To investigate whether the molecular consequences of *sqd-1* depletion in *adr-2(−)* animals align with the phenotypic rescue, RNA sequencing was performed on young adult wild-type and *adr-2(−)* animals treated with either control or *sqd-1* RNAi. First, to determine how *sqd-1* affects gene expression, sequencing reads were aligned to the *C. elegans* reference genome and differential gene expression analysis was performed using DESeq2 (Love and others 2014). Compared to animals treated with control RNAi, *sqd-1* RNAi resulted in significantly altered expression of 8501 transcripts, including 4828 upregulated and 3673 downregulated genes (**Figure 7A, Table S7**). Consistent with the masculinization of *sqd-1* RNAi germlines, the transcripts significantly upregulated by *sqd-1* RNAi significantly overlapped with a previously published dataset of spermatogenic transcripts (Ortiz and others 2014). Furthermore, consistent with the lack of oocytes in the *sqd-1* RNAi germlines, the transcripts significantly downregulated by *sqd-1* RNAi significantly overlapped with oogenic transcripts (Ortiz and others 2014) (**Figure 7B**). Together, these data demonstrate that depletion of *sqd-1* masculinizes the hermaphrodite germline and disrupts the switch from spermatogenesis to oogenesis.

**Figure 7:**
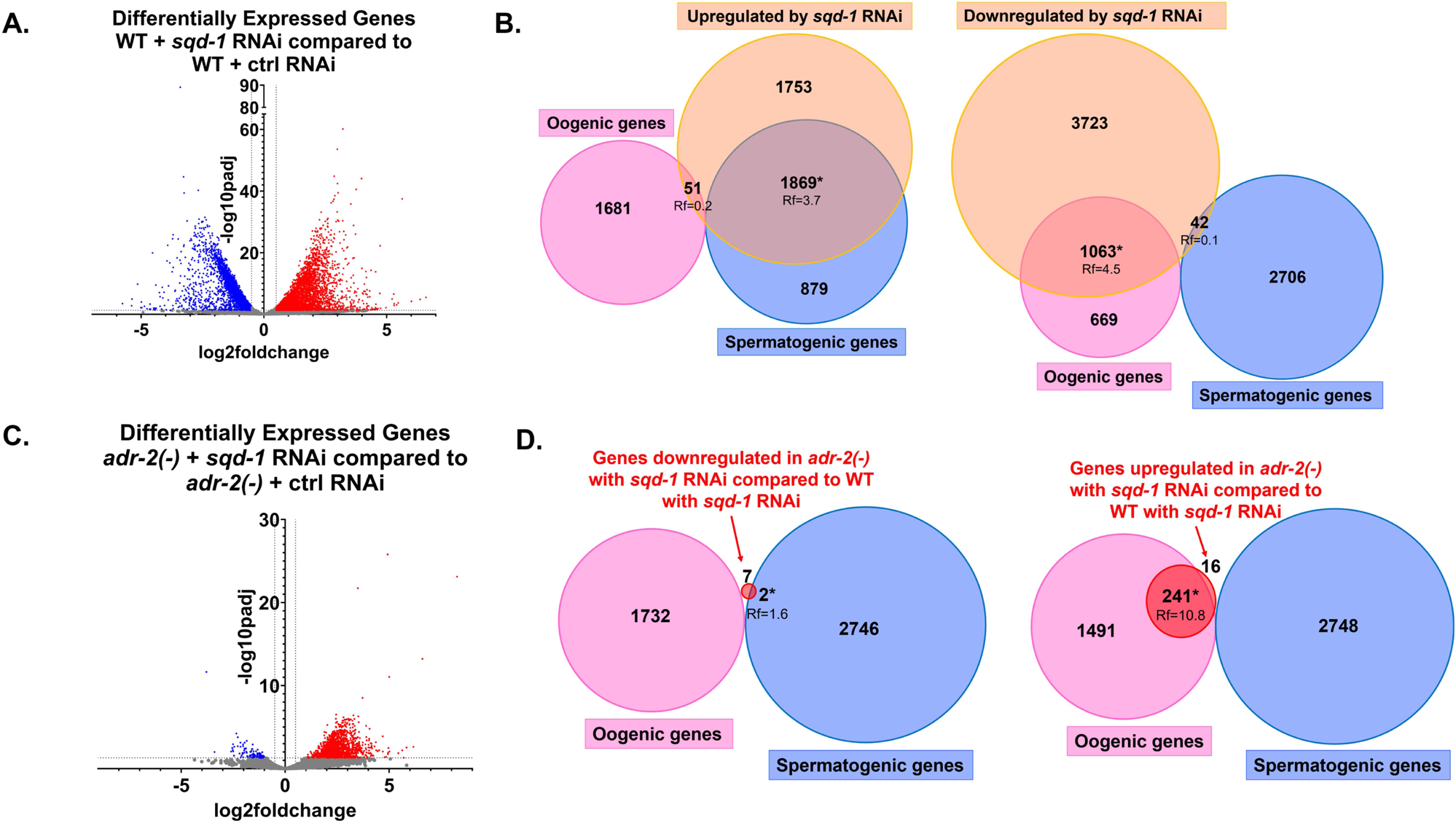
Germline gene expression is consistent with morphology for wild-type and *adr-2(−)* animals depleted of *sqd-1*. A) Differential gene expression of wild-type animals treated with *sqd-1* RNAi compared to wild-type animals treated with control RNAi (data in **Table S7**) Red points indicate significantly upregulated genes (padj <0.05), blue points indicate significantly downregulated genes (padj <0.05), gray points indicate genes that were not significantly differentially expressed. B) Overlap of genes differentially expressed in wild-type animals treated with *sqd-1* RNAi compared to wild-type animals treated with control RNAi (**Table S7**) with spermatogenic and oogenic gene datasets (Ortiz and others 2014). Rf = representation factor, * indicates significant overlap (Rf ≥ 1). C) Differential gene expression of *adr-2(−)* animals treated with *sqd-1* RNAi compared to *adr-2(−)* animals treated with control RNAi (data in **Table S9**). Red points indicate significantly upregulated genes (padj <0.05), blue points indicate significantly downregulated genes (padj <0.05), gray points indicate genes that were not significantly differentially expressed. D) Overlap of genes differentially expressed in *adr-2(−)* animals treated with *sqd-1* RNAi compared to wild-type animals treated with *sqd-1* RNAi (**Table S10**) with spermatogenic and oogenic gene datasets (Ortiz and others 2014). Rf = representation factor, * indicates significant overlap (Rf ≥ 1).

Compared to wild-type animals treated with control RNAi, *adr-2(−)* animals treated with control RNAi had significantly altered expression (padj ≤ 0.05) of only *adr-2* and no other genes (**Figure S6, Table S8**), suggesting that loss of *adr-2* alone does not significantly affect the gene expression of any transcripts in young adult *C. elegans* at the whole worm level. In contrast, when *sqd-1* was depleted from *adr-2(−)* animals, 1408 genes were misregulated compared to *adr-2(−)* animals treated with control RNAi (**Figure 7C, Table S9**). These data suggest that loss of *sqd-1* and *adr-2* results in more alterations in gene expression than loss of *adr-2* alone. However, as depletion of *sqd-1* resulted in over 8000 misregulated transcripts, the vast majority of altered gene expression that occurs upon depletion of *sqd-1* is corrected upon loss of *adr-2*. This latter point likely reflects the comparatively less severe oogenesis defects observed in *adr-2(−)* animals treated with *sqd-1* RNAi (**Figures 5D, 6B**). Consistent with the increase in oogenesis (**Figures 5D, 6B**), transcripts significantly downregulated (padj ≤ 0.05) in *adr-2(−)* animals treated with *sqd-1* RNAi compared to wild-type animals treated with *sqd-1* RNAi (**Table S10**) overlapped significantly (representation factor = 1.6) with spermatogenic transcripts (Ortiz and others 2014), and transcripts significantly upregulated (padj ≤ 0.05) (**Table S10**) overlapped significantly (representation factor = 10.8) with oogenic transcripts (Ortiz and others 2014) (**Figure 7D**). Thus, the changes in gene expression between *adr-2(−)* animals treated with *sqd-1* RNAi and wild-type animals treated with *sqd-1* RNAi reflect the phenotypic shift toward oogenesis. Importantly, however, the loss of *adr-2* does not fully restore oogenic gene expression or abrogate spermatogenic gene expression in animals treated with *sqd-1* RNAi (**Figures 7A and C**), which is expected given that loss of *adr-2* only partially restores fertility in these animals (**Figure 3C**). Together, these data suggest that ADR-2 negatively regulates the switch from spermatogenesis to oogenesis in the absence of *sqd-1*.

### ADR-2 is expressed in nuclei of oogenic germ cells, but not in sperm

To get a better idea of how ADR-2 may be impacting fertility in the absence of *sqd-1*, immunostaining was performed on dissected germlines. First, ADR-2 expression was analyzed in wild-type animals that contained a N-terminal FLAG epitope-tag engineered into the *adr-2* locus. These worms have wildtype ADR-2 function (Dhakal and others 2024). ADR-2 expression was observed in the nuclei of germ cells in oogenic young adult germlines (**Figure 8A**), while an absence of ADR-2 staining is observed in the nuclei of sperm in the spermatheca. Our colleagues have reported similar expression observed with immunostaining using a custom ADR-2 antibody (Eliad and others 2023). In spermatogenic L4 germlines, we also see nuclear expression of ADR-2 in undifferentiated germ cells and a lack of ADR-2 staining in sperm nuclei (**Figure 8B**). Of note, ADR-2 appears to be expressed in the same cells as SQD-1, but in a different subcellular region (**Figure 8C**). Thus, it is likely that the two indirectly interact to regulate oogenesis.

**Figure 8:**
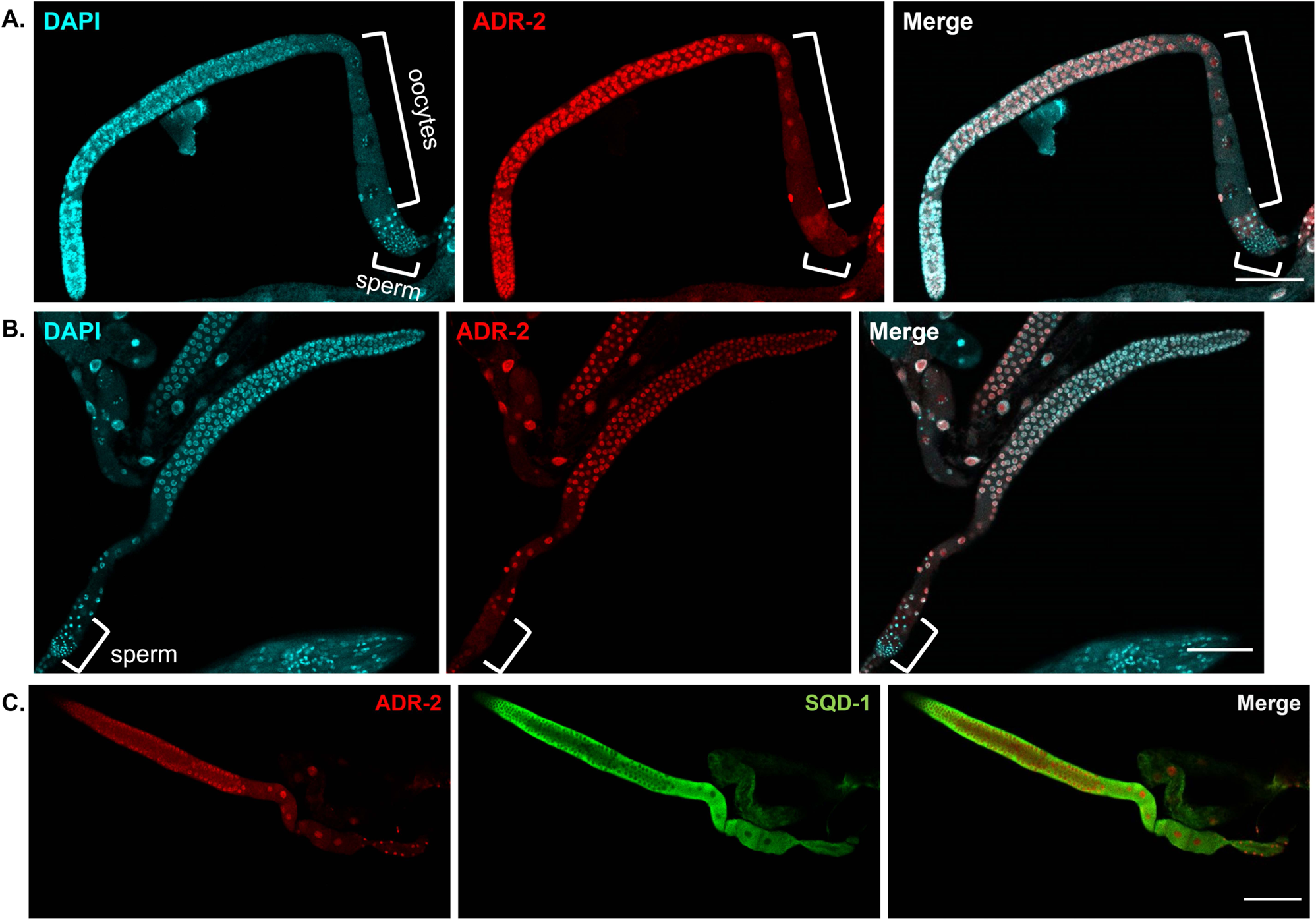
ADR-2 and SQD-1 germline expression patterns. A) Representative confocal images of dissected germlines from young adult (oogenic) animals stained for DAPI (cyan) and ADR-2 (red). Scale bars = 50 μm. Brackets denote oocytes and sperm. B) Representative confocal images of dissected germlines from L4 (spermatogenic) animals stained for DAPI (cyan) and ADR-2 (red). Scale bars = 50 μm. Brackets denote sperm. C) Representative confocal images of dissected germlines from young adult animals stained for ADR-2 (red) and SQD-1 (green). Scale bars= 50μm.

It is possible that loss of SQD-1 could result in mislocalization of ADR-2, which may lead to the oogenesis defects. To explore this possibility, ADR-2 staining using a custom ADR-2 antibody (Rajendren and others 2018) was performed on *sqd-1* null germlines. Similar to wild-type animals, ADR-2 staining is observed in the nuclei of undifferentiated germ cells, but not in sperm nuclei (**Figure S7**), suggesting ADR-2 localization is unaltered in the absence of *sqd-1*. This finding further indicates that the mechanism of ADR-2 impacting fertility in the absence of *sqd-1* is likely not via a direct protein-protein interaction. As ADR-2 and SQD-1 are both RBPs, it is possible that ADR-2 and SQD-1 share a common target RNA. We hypothesize that ADR-2 binding to the target RNA in wild-type animals does not impact fertility due to additional downstream regulation by SQD-1 (**Figure 9B**). However, it is also possible that ADR-2 binds a new target that is only expressed in the absence of *sqd-1* (**Figure 9B**).

**Figure 9:**
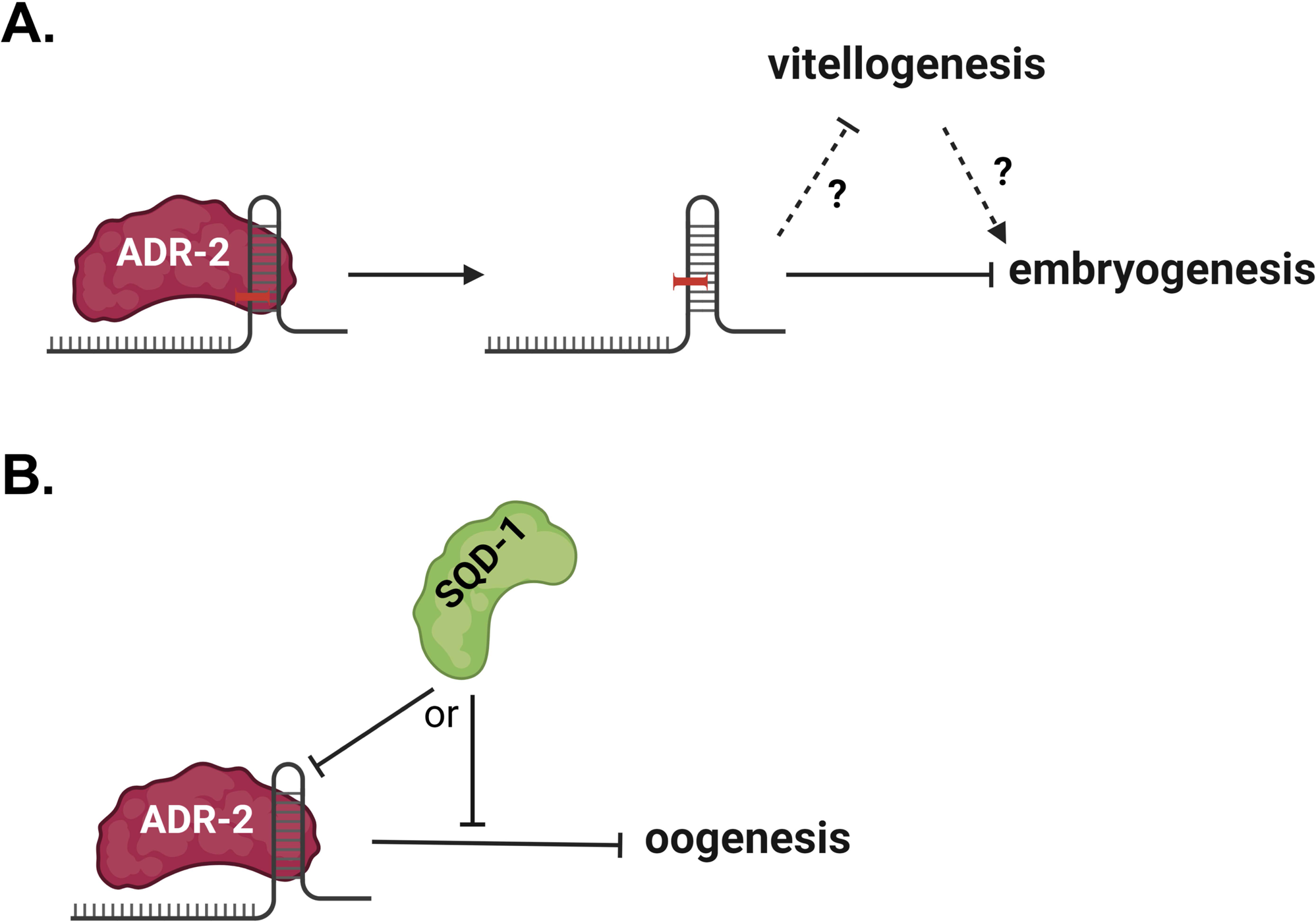
A) Model for the role of A-to-I editing by ADR-2 in regulating embryogenesis and vitellogenesis in the *vit-2::gfp* background. B) Model for the role of ADR-2 in regulating oogenesis along with SQD-1. Figures created with BioRender.

## Discussion

RBPs are known to play important roles in regulating and facilitating reproduction in all organisms (Albarqi and Ryder 2022; Nguyen-Chi and Morello 2011; Rosario and others 2017). From the results of this study, we feel that ADARs can be added to this list. In this study, we present evidence from two genetic backgrounds that loss of *adr-2* modulates reproductive function in *C. elegans*. First, we show that loss of editing by ADR-2 restores wild-type fertility in subfertile *vit-2::gfp* animals (**Figure 1**), suggesting that editing by *adr-2* suppresses fertility in the *vit-2::gfp* background (**Figure 9A**). Second, we show that loss of *adr-2* partially rescues sterility in animals treated with *sqd-1* RNAi by rescuing oogenesis (**Figures 3C and 6**), suggesting that ADR-2 acts to suppress oogenesis in the absence of *sqd-1* (**Figure 9B**). These examples suggest a potential broader role for *adr-2* in suppressing fertility in certain mutant backgrounds. Previous studies have observed positive effects of ADARs on fertility in *C. elegans*, with decreased fertility reported upon loss of both ADARs (*adr-1* and *adr-2*) in combination with factors from small RNA pathways (Fischer and Ruvkun 2020; Reich and others 2018). Our work is the first to indicate that ADARs can negatively impact fertility, and the first to show *adr-1*-independent effects of *adr-2* on fertility. In future studies, it would be of interest to assess the effects of ADARs on germline RNA regulation more globally to better understand how they regulate reproductive processes. Testis-specific roles for deaminase-deficient ADAR family members, ADADs, have previously been identified (Islam and others 2023; Snyder and others 2020), and A-to-I editing has been reported in human ovaries (Tan and others 2017). Profiling A-to-I editing in the *C. elegans* germline would provide clues as to the roles of ADARs in maintaining proper reproductive function. Furthermore, while we reported almost no effects of loss of *adr-2* on RNA expression when analyzed at the organismal level (**Figure S6, Table S8**), our prior work has suggested that ADARs play important roles in tissue-specific gene regulation (Deffit and others 2017; Mahapatra and others 2023) and thus, future work should focus on assessing the effects of loss of ADARs specifically on the germline transcriptome.

We show here that ADR-2 is expressed in undifferentiated germ nuclei and in oocyte nuclei (**Figure 8**), which points to possible roles for ADARs in these cells. However, it is important to note that, due to the interplay between the germline and other tissues (Devanapally and others 2015; Gopal and others 2021; Qi and others 2021; Starich and others 2020), the role of ADR-2 in suppressing fertility in *vit-2::gfp* and *sqd-1*-depleted animals may arise from expression/functions in a non-germline tissue. Given that vitellogenin is produced in the intestine, it is possible that *adr-2* regulates vitellogenin synthesis in the intestine, and that the fertility defects in *vit-2::gfp* animals are a downstream effect of intestinal ADR-2 expression. As the *vit-2::gfp* phenotype is editing-dependent, assessment of ADR-2 editing targets in *vit-2::gfp* animals compared to wild-type animals may help to uncover the molecular mechanism behind this fertility defect.

Using *adr-2(−);vit-2::gfp* animals as a reporter, a high-throughput RNAi screen was performed and identified 23 RBPs that regulate embryo production and/or yolk provisioning in *C. elegans*. While some of these are known regulators of germline development, many do not have well-described roles in yolk provisioning or embryogenesis (**Table S5**). Of the 23 RBPs, 11 have characterized molecular functions in oogenesis or embryogenesis that likely contribute to the fertility phenotypes identified in the screen (**Table S5**). The 12 remaining RBPs have come up in previous screens for factors affecting germline morphology (Green and others 2011; Updike and Strome 2009), germline differentiation and sex determination (Kalis and others 2010; Kerins and others 2010; Wang and others 2012), RME (Balklava and others 2007), as well as general RNAi-based phenotypic screens (Kamath and others 2003; Maeda and others 2001); however specific roles of these RBPs in embryogenesis are not yet described. We suggest that the *adr-2(−);vit-2::gfp* strain could be used in future studies as a method for interrogating the function of mutations within the RBPs. Additionally, as we showed that the *vit-2::gfp* fertility defect is dependent on RNA editing activity by ADR-2 (**Figure 1E**), we suggest that this background may be useful in future studies to identify regulators of RNA editing activity by ADR-2.

One interesting observation from our study was that, while unable to restore fertility (measured as embryo counts) in animals treated with RNAi against *csr-1*, *cgh-1*, and *sfa-1*, loss of *adr-2* resulted in increased vitellogenin accumulation in the body cavity (**Figure 3A**), suggesting *adr-2* may act to suppress vitellogenesis or promote proper vitellogenin provisioning in certain backgrounds. This regulation of vitellogenesis or vitellogenin provisioning may contribute to the brood size defects observed in *vit-2::gfp* animals (**Figure 9A**). The type of vitellogenin accumulation seen in *adr-2(−);vit-2::gfp* animals treated with RNAi against *csr-1*, *cgh-1*, *sfa-1*, and *sqd-1* has previously been seen in animals with defects in RME, the process by which yolk proteins are imported into developing oocytes (Balklava and others 2007; Balklava and others 2016; Grant and Hirsh 1999). Depletion of *cgh-1* and *sfa-1* have previously been associated with defects in RME (Balklava and others 2007), and it is likely that RME is disrupted when *sqd-1* or *csr-1* is absent as well. Previous studies have suggested that vitellogenin production is normally subject to regulation by signals from the germline, ensuring there is enough vitellogenin to load into the oocytes being produced (Balklava and others 2016; DePina and others 2011). However, if this communication is somehow altered, vitellogenin production could become uncoupled to oocyte production, leading to accumulation of vitellogenin in the body cavity (Balklava and others 2016). Thus, it is possible that *cgh-1*, *csr-1*, *sqd-1*, and *sfa-1* may play roles in the coupling of vitellogenin production to oocyte production, which would account for the accumulation of vitellogenin when these factors are lost. Future experiments, likely using other methods or reporters to assess vitellogenin provisioning, are needed to investigate whether these factors affect these processes in other backgrounds, and to assess the effects of each factor on vitellogenin provisioning in backgrounds with altered oogenesis.

This work also provides the first characterization of the effects of SQD-1 on germline morphology and gene expression in *C. elegans*. We show that loss of *sqd-1* leads to masculinization of the germline, characterized by a failure to switch from spermatogenesis to oogenesis in early adulthood (**Figure 5A-D**). Additionally, we show that these functional defects are associated with the misexpression of thousands of genes, including downregulation of oogenic genes and upregulation of spermatogenic genes (**Figure 7A-B**). We show that loss of *adr-2* is sufficient to rescue the switch from spermatogenesis to oogenesis, however this rescue is not sufficient to fully restore fertility to wild-type levels (**Figure 6**, **Figure 3C**). While loss of *adr-2* rescues the expression of many of the genes that are misexpressed due to *sqd-1* depletion, there are also a large number of genes that are still misexpressed in *adr-2(−)* animals treated with *sqd-1* RNAi compared to animals treated with control RNAi (**Figure 7C, Tables S7,S9**). Thus, it is possible that the SQD-1 regulated genes that are rescued by loss of *adr-2* contribute to the sperm to oocyte switch, while the genes that are not rescued by loss of *adr-2* contribute to the overall fertility. It would be useful in uncovering the molecular mechanism behind the editing-independent infertility in *sqd-1* depleted animals to assess ADR-2 binding targets in the presence and absence of *sqd-1*.

While further research is needed to elucidate the molecular mechanism by which *sqd-1* facilitates the switch from spermatogenesis to oogenesis in wild-type animals, the known roles of its homologs in other species may provide clues. In *Drosophila*, *squid* mutant embryo defects stem from the mislocalization of *gurken* mRNA; Gurken is an EGFR ligand whose dorsal-specific localization during later stages of oogenesis is necessary for proper embryo patterning (Norvell and others 1999). Similarly, it is possible that the sterility observed in *C. elegans* depleted of *sqd-1* could be a result of mislocalization of a specific mRNA target of SQD-1, which in turn leads to the misexpression of several downstream genes. Additionally, *Drosophila squid* has been shown to associate with other factors to coordinate translational repression of specific transcripts important in embryogenesis (Clouse and others 2008). It is possible *C. elegans* SQD-1 could play a similar role in coordinating translational repression, as SQD-1 has been identified in complex with GLD-1, a known translational repressor (Akay and others 2013). In future studies, identification of the RNA targets of SQD-1 will help elucidate its role in the well-defined germline sex determination pathway (Ellis and Schedl 2007). Tools developed in our study, particularly the V5-tagged SQD-1 strain, as well as data generated here, such as SQD-1 localization and the transcriptomic changes upon loss of *sqd-1*, could be useful in these future studies.

## Data availability

Strains and plasmids are available upon request. Raw and processed high-throughput sequencing data generated in this study have been submitted to the NCBI Gene Expression Omnibus (GEO; https://www.ncbi.nlm.nih.gov/geo/) under accession number GSE244602.

## Acknowledgements

The authors would like to acknowledge Christiane Hassel and the Indiana University Bloomington Flow Cytometry Core Facility for the use and operation of the COPAS SELECT, the Indiana University Center for Genomics and Bioinformatics for running the high-throughput sequencing experiments, Dr. Scott Aoki (Indiana University School of Medicine) for his assistance with learning gonad extrusion and staining and his feedback on this manuscript, the Indiana University Light Microscopy Imaging Center for the use of the Leia SP8 Confocal, and Dr. Lesley Weaver (IU Biology) for her feedback on the micrographs presented in the manuscript. The authors would also like to acknowledge Dr. Dustin Updike of Mount Desert Island Biological laboratory as well as members of the Hundley lab: Dr. Chinnu Salim, Ananya Mahapatra, Boyoon Yang, Alfa Dhakal, and Emma Lamb for their feedback and assistance on this manuscript.

## Funding

This work was supported in part by the National Institutes of Health/National Institute of General Medical Sciences [R01GM130759] awarded to H.A.H, including supplemental funding for equipment [R01GM130759-03S1], the National Institutes of Health/National Institute of General Medical Sciences [T32 GM131994] and the National Institutes of Health/National Institute of Child Health and Human Development [F31 HD110244-01] awarded to E.E., and the National Center for Advancing Translational Sciences, Clinical and Translational Sciences Award (Indiana University Medical Student Program for Research and Scholarship (IMPRS) to M.B.). Some strains were provided by the *Caenorhabditis* Genetics Center (CGC), which is funded by NIH Office of Research Infrastructure Programs [P40 OD010440].

## Conflict of Interest

The authors declare no conflict of interest.

## Author contributions

Designed the experiments: E.E., H.A.H.

Created reagents: S.P. (HAH56 strain), E.E. (HAH51-56, 63-64 strains)

Performed the experiments: E.E., M.F. (secondary screen), M.B. (primary screen)

Performed the bioinformatics analysis: E.E.

Wrote and edited the manuscript: E.E., H.A.H.

---

**Figure S1:** A second *adr-2* mutant allele restores embryo count in *vit-2::gfp* animals. Embryo content of the indicated strains. Data represents 3 biological replicates, n=20 worms/replicate. Statistical significance determined via ordinary one-way Anova; ns= not significant (p>0.05), * indicates p≤0.05. Note: this assay was performed at an earlier timepoint than other embryo counting assays, resulting in overall reduced embryo contents.

**Figure S2:** Loss of *adr-1* does not restore embryo count in *vit-2::gfp* animals. Embryo content of the indicated strains. Data represents 3 biological replicates, n=20 worms/replicate. Statistical significance determined via ordinary one-way Anova; ns = not significant (p>0.05).

**Figure S3:** A) Primary screen results. *adr-2(−);vit-2::gfp* animals were fed with RNAi against 255 RNA binding proteins. Animals were collected and average GFP fluorescence in each population was measured using the COPAS select. Analysis was restricted to worms meeting length and density criteria for adult worms. Treatments causing no significant change (gray points), >2.5 fold increase (red points), and >2.5 fold decrease (blue points) in GFP fluorescence compared to positive control (empty vector RNAi) treated animals are plotted. n=2000-5000 animals per treatment. B) RNAi vectors that caused significant change in GFP levels in the primary screen (A) were tested a second time to eliminate unreproducible results (n=2000-5000 animals per replicate). Solid points indicate average fluorescence values for each RNAi treatment in primary screening replicate (red= increased compared to control, blue= decreased compared to control). Open symbols indicate average fluorescence values for each RNAi treatment in second screening replicate (gray= no significant difference compared to control, red= increased compared to control, blue= decreased compared to control). Treatments whose second replicates showed no significant difference from controls (*psf-1*, ZK686.2) or the opposite patten from the primary replicate (*nuo-2*, F55F8.2, *rbd-1*) were not designated high-confidence modulators of GFP level in *adr-2(−);vit-2::gfp* animals.

**Figure S4:** *sqd-1* RNAi efficiency is unaffected in *adr-2(−)* animals compared to wild-type. qPCR was performed to measure *sqd-1* mRNA in wild-type animals treated with control RNAi and *sqd-1* RNAi and *adr-2(−)* animals treated with control RNAi and *sqd-1* RNAi. Plotted expression values are normalized to *gpd-3* (housekeeping gene) expression. Data represents 3 biological replicates. Significance determined via unpaired t-test, ns indicates no significant difference (p>0.05).

**Figure S5:** Representative confocal images of dissected germlines from day 2 adult (96 hours post egg-lay) animals stained with DAPI. Top panel = wild-type, bottom panel = *sqd-1* null. Scale bars = 50 μm.

**Figure S6:** Differential gene expression of *adr-2(−)* animals treated with control RNAi compared to wild-type animals treated with control RNAi (data in **Table S8**). Gray points indicate genes that were not significantly differentially expressed (padj > 0.05).

**Figure S7:** Representative confocal images of dissected germlines from A) day 1 adult (72 hours post egg-lay) and B) day 2 adult (96 hours post egg-lay) *sqd-1* null animals stained for DAPI and ADR-2. Brackets denote sperm. Scale bars = 50 μm.

## Supplementary Tables

**Table S1:** Oligonucleotides used in this study.

**Table S2:** RNAi feeding vectors used in screen for regulators of fertility. Related to **Figures 2, 3, 5-7, S3-4, S6**. Column A indicates RNAi vector number used in this study, Column B indicates the name of the gene targeted by each RNAi vector, Column C indicates the corresponding WormbaseID, Column D indicates the type and number of RNA binding domains in each gene. Column E contains a description of each RNAi feeding vector, and Column F contains the references for the RNA binding domain information in Column D. References: (Mistry and others 2021; Paysan-Lafosse and others 2023; Sigrist and others 2013; Tamburino and others 2013)

**Table S3:** Primary screen data-*adr-2(−);vit-2::gfp* animals (Related to **Figures S3** and **2C,D**). Column A indicates RNAi vector ID as listed in **Table S2**. Column B indicates the gene target of each RNAi vector, Column C indicates the corresponding WormbaseID. Column D indicates the average Time of Flight (TOF) value measured by the COPAS SELECT, Column E indicates the average extinction (ext) value measured by the COPAS SELECT, and Column F indicates the ratio of average TOF over ext. Column G indicates the average GFP fluorescence measured by the COPAS SELECT, and Column H indicates the number of animals analyzed in each experiment. Columns J, K, and L indicate normalized values for GFP, TOF, and ext, respectively. Normalized values are calculated by dividing the average value for each RNAi vector by the average value for the control vector in each experiment. Normalized GFP values highlighted green indicate ≥2.5-fold decrease in GFP compared to control, values highlighted red indicate ≥ 2.5-fold increase in GFP compared to control.

**Table S4:** Secondary screen data (Related to **Figures S3** and **2C,D**). First page contains data for *adr-2(−);vit-2::gfp* animals, second page contains data for *vit-2::gfp* animals. Column A indicates RNAi vector ID as listed in **Table S2**. Column B indicates the gene target of each RNAi vector, Column C indicates the corresponding WormbaseID, Column D indicates the average Time of Flight (TOF) value measured by the COPAS SELECT, Column E indicates the average extinction (ext) value measured by the COPAS SELECT, and Column F indicates the ratio of average TOF over ext. Column G indicates the average GFP fluorescence measured by the COPAS SELECT, and Column H indicates the number of animals analyzed in each experiment. Columns J, K, and L indicate normalized values for GFP, TOF, and ext, respectively. Normalized values are calculated by dividing the average value for each RNAi vector by the average value for the control vector in each experiment. Normalized GFP values highlighted green indicate ≥2.5-fold decrease in GFP compared to control, values highlighted red indicate ≥ 2.5-fold increase in GFP compared to control, values highlighted yellow indicate fold changes ≤2.5. On first page, column N indicates the normalized GFP value from the primary screen (**Table S3**).

**Table S5:** Previously described germline roles and genome-wide screen hits for RBPs designated high-confidence modulators of GFP level in *adr-2(−);vit-2::gfp* animals. Related to **Figures 2-3, S3**. References: (Balklava and others 2007; Barbee and Evans 2006; Barbee and others 2002; Elbaum-Garfinkle and others 2015; Francis and others 1995; Fujita and others 1998; Graham and others 1993; Green and others 2011; Hubert and Anderson 2009; Kalis and others 2010; Kamath and others 2003; Kerins and others 2010; Ko and others 2013; Maeda and others 2001; Navarro and others 2001; Singh and others 2017; Subramaniam and Seydoux 2003; Updike and Strome 2009; Wang and others 2012; Wedeles and others 2013)

**Table S6:** Quality measures for high-throughput sequencing. Related to **Figure 7**. Column A lists sequenced samples. Total Sequences (Column B) and Sequences Flagged as Poor Quality (Column C) determined via FASTQC. % uniquely mapped reads (Column D) determined via STAR alignment.

**Table S7:** Differentially expressed genes in wild-type animals treated with *sqd-1* RNAi compared to wild-type animals treated with control RNAi (Related to **Figure 7**). First page contains all differentially expressed genes Padj ≤ 0.05. Second page contains genes downregulated in wild-type animals treated with *sqd-1* RNAi compared to wild-type animals treated with control RNAi (Padj ≤0.05). Third page contains genes upregulated in wild-type animals treated with *sqd-1* RNAi compared to wild-type animals treated with control RNAi (Padj ≤0.05). For each page, Column A indicates WormbaseID of each gene, Column B indicates corresponding gene name, Column C indicates log_2_fold change value from DESeq, Column D indicates pvalue from DESeq, Column E indicates padj from DESeq, Columns F-H indicate expression values from DESeq for each of three replicates of wild-type animals treated with *sqd-1* RNAi, Columns I-K indicate expression values from DESeq for each of three replicates of wild-type animals treated with control RNAi.

**Table S8:** Differentially expressed genes in *adr-2(−)* animals treated with control RNAi compared to wild-type animals treated with control RNAi (Related to **Figure S6**). Column A indicates WormbaseID of each gene, Column B indicates corresponding gene name, Column C indicates log_2_fold change value from DESeq, Column D indicates pvalue from DESeq, Column E indicates padj from DESeq, Columns F-H indicate expression values from DESeq for each of three replicates of *adr-2(−)* animals treated with control RNAi, Columns I-K indicate expression values from DESeq for each of three replicates of wild-type animals treated with control RNAi.

**Table S9:** Differentially expressed genes in *adr-2(−)* animals treated with *sqd-1* RNAi compared to *adr-2(−)* animals treated with control RNAi (Related to **Figure 7**). First page contains all differentially expressed genes Padj ≤ 0.05. Second page contains genes downregulated in *adr-2(−)* animals treated with *sqd-1* RNAi compared to *adr-2(−)* animals treated with control RNAi (Padj ≤0.05). Third page contains genes upregulated in *adr-2(−)* animals treated with *sqd-1* RNAi compared to *adr-2(−)* animals treated with control RNAi (Padj ≤0.05). For each page, Column A indicates WormbaseID of each gene, Column B indicates corresponding gene name, Column C indicates log_2_fold change value from DESeq, Column D indicates pvalue from DESeq, Column E indicates padj from DESeq, Columns F-H indicate expression values from DESeq for each of three replicates of *adr-2(−)* animals treated with *sqd-1* RNAi, Columns I-K indicate expression values from DESeq for each of three replicates of *adr-2(−)* animals treated with control RNAi.

**Table S10:** Differentially expressed genes in *adr-2(−)* animals treated with *sqd-1* RNAi compared to wild-type animals treated with *sqd-1* RNAi (Related to **Figure 7D**). First page contains all differentially expressed genes Padj ≤ 0.05. Second page contains genes downregulated in *adr-2(−)* animals treated with *sqd-1* RNAi compared to wild-type animals treated with *sqd-1* RNAi (Padj ≤0.05). Third page contains genes upregulated in *adr-2(−)* animals treated with *sqd-1* RNAi compared to wild-type animals treated with *sqd-1* RNAi (Padj ≤0.05). For each page, Column A indicates WormbaseID of each gene, Column B indicates corresponding gene name, Column C indicates log_2_fold change value from DESeq, Column D indicates pvalue from DESeq, Column E indicates padj from DESeq, Columns F-H indicate expression values from DESeq for each of three replicates of *adr-2(−)* animals treated with *sqd-1* RNAi, Columns I-K indicate expression values from DESeq for each of three replicates of wild-type animals treated with *sqd-1* RNAi.

## Notes

### Competing Interest Statement

The authors have declared no competing interest.

### Summary of Updates

New experimental results demonstrating ADR-2 and SQD-1 localization in the germline. Additional data on the loss of sqd-1 phenotype.

